# BRCA1/BRC-1 and SMC-5/6 regulate DNA repair pathway engagement during *C. elegans* meiosis

**DOI:** 10.1101/2022.06.12.495837

**Authors:** Erik Toraason, Alina Salagean, David E. Almanzar, Ofer Rog, Diana E. Libuda

## Abstract

The preservation of genome integrity during sperm and egg development is vital for reproductive success. During meiosis, the tumor suppressor BRCA1/BRC-1 and structural maintenance of chromosomes 5/6 (SMC-5/6) complex genetically interact to promote high fidelity DNA double strand break (DSB) repair, but the specific DSB repair outcomes these proteins regulate remain unknown. Here we show that BRCA1/BRC-1 and the SMC-5/6 complex limit intersister crossover recombination as well as error-prone repair pathways during meiotic prophase I. Using genetic and cytological methods to monitor repair of DSBs with different repair partners in *Caenorhabditis elegans*, we demonstrate that both BRC-1 and SMC-5/6 repress intersister crossover recombination events, with meiotic cells becoming more dependent upon these proteins to repair DSBs in late meiotic prophase I. Sequencing of conversion tracts from homolog-independent DSB repair events indicates that BRC-1 regulates intersister/intrachromatid noncrossover conversion tract length. Moreover, we find that BRC-1 also specifically inhibits error prone repair of DSBs induced at mid-pachytene. Finally, we reveal that functional BRC-1 enhances DSB repair defects in *smc-5* mutants by repressing theta-mediated end joining (TMEJ). Taken together, our study illuminates the coordinate interplay of BRC-1 and SMC-5/6 to regulate DSB repair outcomes in the germline.

## Introduction

Meiosis is the specialized form of cell division by which most sexually reproducing organisms generate haploid gametes. In a diploid organism, each meiotic cell begins prophase I with four copies of the genome – two homologous chromosomes (homologs) and an identical replicate of each homolog called a sister chromatid. As mutations incurred in the gamete genome will be passed on to the resultant progeny, it is crucial that genome integrity be maintained during meiosis. Despite this risk, a highly conserved feature of the meiotic program is induction of DNA double strand breaks (DSBs) by the topoisomerase-like protein Spo11 (Keeney *et al*. 1997). A limited subset of DSBs must engage the homologous chromosome as a recombination partner and be resolved as a crossover event, which forges a physical connection between homologs that facilitates accurate chromosome segregation at the meiosis I division. DSBs are incurred in excess of the number of eventual crossovers, therefore other pathways must be utilized to repair residual DSBs. How meiotic cells regulate repair pathway engagement to both accurately and efficiently resolve DSBs is a critical question in the field of genome integrity.

The majority of meiotic DSBs are repaired through interhomolog noncrossover recombination mechanisms (Hunter 2015). Multiple models are proposed for how meiotic noncrossover repair occurs. Evidence in *Drosophila* suggests that both interhomolog noncrossovers and crossovers may be generated by differential processing of similar joint molecule intermediates (Crown *et al*. 2014). Work in budding yeast, mammals, and *Arabidopsis* indicates that the majority of interhomolog noncrossovers are generated via synthesis-dependent strand annealing (SDSA) with the homolog (Hunter 2015). In SDSA, the resected end of the DSB invades a repair template, synthesizes new sequence, dissociates from its repair template, and finally utilizes the synthesized sequence to anneal to the other resected end of the DSB.

Meiotic DSBs may also be resolved by recombination with the sister chromatid (Schwacha and Kleckner 1997; Goldfarb and Lichten 2010; Toraason *et al*. 2021a; Almanzar *et al*. 2021). In budding yeast, DSB resolution by intersister recombination is disfavored so as to promote DSB repair with the homologous chromosome in wild type conditions (Schwacha and Kleckner 1994, 1997; Goldfarb and Lichten 2010; Kim *et al*. 2010; Humphryes and Hochwagen 2014). In metazoan meiosis, however, the engagement of intersister repair has proven challenging to detect and quantify. While recombination between polymorphic homologs may be readily studied via sequence conversions in final repair products, the identical sequences of sister chromatids preclude the detection of intersister recombination by sequencing-based approaches. Recently, two methods have been developed in the nematode *Caenorhabditis elegans* to enable direct detection of homolog-independent meiotic recombination (Toraason *et al*. 2021a; Almanzar *et al*. 2021). Toraason *et al*. 2021a constructed an intersister/intrachromatid repair (ICR) assay, which exploits nonallelic recombination at a known locus in the genome to identify homolog-independent repair events in resultant progeny. Almanzar *et al*. 2021 designed an EdU labeling assay to cytologically identify sister chromatid exchanges (SCEs) in compacted chromosomes at diakinesis. Together, these studies demonstrated that: 1) homolog-independent meiotic recombination occurs in *C. elegans*; 2) the sister chromatid and/or same DNA molecule is the exclusive recombination repair template in late prophase I; and, 3) intersister crossovers are rare and represent a minority of homolog-independent recombination products (Toraason *et al*. 2021a; Almanzar *et al*. 2021).

While meiotic cells primarily utilize recombination to resolve DSBs, error prone repair pathways are also available in meiosis to repair DSBs at the risk of introducing *de novo* mutations (Gartner and Engebrecht 2022). These error prone mechanisms are repressed to promote recombination repair, but are activated in mutants that disrupt recombination (Lemmens *et al*. 2013; Yin and Smolikove 2013; Macaisne *et al*. 2018; Kamp *et al*. 2020). Non-homologous end joining (NHEJ), which facilitates the ligation of blunt DNA ends by the DNA ligase IV homolog LIG-4, is active in the *C. elegans* germline (Yin and Smolikove 2013; Macaisne *et al*. 2018). Recent studies have indicated that microhomology-mediated end-joining facilitated by the DNA polymerase θ homolog POLQ-1 (theta-mediated end-joining, TMEJ) is the primary pathway by which small mutations are incurred in *C. elegans* germ cells (Van Schendel *et al*. 2015; Kamp *et al*. 2020). Neither NHEJ nor TMEJ are required for successful meiosis (Colaiácovo *et al*. 2003; Lemmens *et al*. 2013; Volkova *et al*. 2020; Kamp *et al*. 2020), indicating recombination is sufficient for meiotic DSB repair and gamete viability under normal conditions.

The structural maintenance of chromosomes 5/6 complex and tumor suppressor BRCA1 (SMC-5/6 and BRC-1 respectively in *C. elegans*) are highly conserved and regulate meiotic DSB repair in *C. elegans* (Bickel *et al*. 2010; Hong *et al*. 2016; Li *et al*. 2018; Kamp *et al*. 2020). The SMC-5/6 complex is vital for preservation of meiotic genome integrity, as *C. elegans* mutants for *smc-5* exhibit a transgenerational sterility phenotype (Bickel *et al*. 2010). Although null mutations of *smc-5, smc-6,* and *brc-1* revealed that they are not required for development nor reproduction in *C. elegans* (Adamo *et al*. 2008; Bickel *et al*. 2010; Li *et al*. 2018), both SMC-5/6 and BRC-1 are required for efficient DSB repair, as *smc-5* and *brc-1* null mutants both display meiotic chromosome fragmentation at diakinesis indicative of unresolved DSBs (Bickel *et al*. 2010). BRC-1 has also been shown to repress error prone DSB repair via NHEJ and TMEJ (Li *et al*. 2020; Kamp *et al*. 2020). Further, SMC-5/6 and BRC-1 may promote genome integrity in part by facilitating efficient recombination, as *smc-5* and *brc-1* mutants exhibit persistent DSBs marked by the recombinase RAD-51 (Boulton *et al*. 2004; Adamo *et al*. 2008; Bickel *et al*. 2010; Kamp *et al*. 2020), suggesting that early recombination steps are delayed in these mutants. BRC-1 further prevents recombination between heterologous templates to promote accurate recombination repair (León-Ortiz *et al*. 2018). Despite these apparent DNA repair defects, interhomolog crossover formation is largely unaffected by *smc-5* and *brc-1* mutations (Adamo *et al*. 2008; Bickel *et al*. 2010; Li *et al*. 2018). Taken together, these data support the hypothesis that SMC-5/6 and BRC-1 may be required for intersister repair in *C. elegans*.

SMC-5/6 and BRC-1 genetically interact to regulate DSB repair. The incidence of unresolved DSBs in *smc-5* and *brc-1* mutants are not additive in the double *smc-5;brc-1* mutant context, which suggests that SMC-5/6 and BRC-1 may share some DSB repair functions (Bickel *et al*. 2010). Other experiments, however, indicate opposing functions for SMC-5/6 and BRC-1, as both the mitotic DNA replication defects in *smc-5* mutants and the synthetic lethality of *smc-5;him-6* (BLM helicase) double mutants are suppressed by *brc-1* mutation (Wolters *et al*. 2014; Hong *et al*. 2016). Crucially, the specific steps of recombination regulated by SMC-5/6 and BRC-1 which intersect to influence DNA repair outcomes remain unknown.

To determine the DSB repair functions of SMC-5/6 and BRC-1 which regulate DNA repair outcomes during *C. elegans* meiosis, we employed a multipronged approach utilizing genetic assays, cytology, sequence analysis of recombinant loci, and functional DSB repair assays in *smc-5* and *brc-1* mutants. We find that SMC-5/6 and BRC-1 function to repress meiotic intersister crossover recombination, and that BRC-1 specifically regulates homolog-independent noncrossover intermediate processing. Through these experiments, we also find that BRC-1 prevents mutagenic DSB repair at the mid-pachytene stage of meiotic prophase I. By assessing germ cell capacity to resolve exogenous DSBs, we demonstrate that meiotic nuclei become more dependent on SMC-5/6 and BRC-1 for DSB repair in late stages of meiotic prophase I. Finally, we reveal that *smc-5* mutant DSB repair defects are enhanced by functional BRC-1, which impedes gamete viability in part by repressing error prone repair pathways. Taken together, our study defines specific functions and interactions of BRC-1 and SMC-5/6 to regulate meiotic DSB repair outcomes across meiotic prophase I.

## Results

### BRC-1 restricts intersister crossovers

To directly assess the functions of BRC-1 in homolog-independent DSB repair, we employed the recently developed intersister/intrachromatid (ICR) assay (Toraason *et al*. 2021a; b). The ICR assay enables: 1) the controlled generation of a single DSB in *C. elegans* meiotic nuclei via heat shock inducible mobilization of a Mos1 transposon (Bessereau *et al*. 2001; Robert and Bessereau 2007); 2) detection of the repair outcome of the induced DSB with the sister chromatid or same DNA molecule by reconstituting GFP fluorescence in resultant progeny; and, 3) delineation of homolog-independent crossover and noncrossover recombination outcomes (Toraason *et al*. 2021a). Since the *C. elegans* germline is organized in a spatial-temporal gradient in which nuclei move progressively through the stages of meiotic prophase I along the distal-proximal axis (Jaramillo-Lambert *et al*. 2007; Rosu *et al*. 2011; Cahoon and Libuda 2021), oocytes at all stages of meiotic prophase I can be affected simultaneously by a specific treatment, such as heat shock or irradiation. Since the rate of meiotic progression in the *C. elegans* germline is known (Jaramillo-Lambert *et al*. 2007; Rosu *et al*. 2011; Cahoon and Libuda 2021), we can score resultant progeny at specific timepoints post heat shock to distinguish oocytes which incurred a Mos1-excision induced DSB at the stages of prophase I when the homologous chromosome is available as a repair partner (the ‘interhomolog window’, leptotene-mid pachytene, 22-58hr post heat shock) from the stages when the homolog is not readily engaged for DSB repair (the ‘non-interhomolog window’, late pachytene-diplotene, 10-22hr post heat shock) (Rosu *et al*. 2011).

We performed the ICR assay in a *brc-1(xoe4)* mutant, which removes the entire *brc-1* coding sequence (Li *et al*. 2018). If BRC-1 is required for efficient intersister repair, then we expected the overall frequency of ICR assay GFP+ progeny to be reduced. Contrary to this hypothesis, we found that GFP+ progeny were elevated at all interhomolog window timepoints and were not reduced within the non-an overall increase in intersister/intrachromatid repair in *brc-1* mutants (see Methods). Regardless of the absolute number of ICR assay GFP+ progeny, we identified both crossover and noncrossover interhomolog window (Supplemental Figure 1A). This result could be explained by multiple effects, such as altered repair template bias, and therefore does not necessarily represent recombinant progeny at all timepoints scored (Supplemental Figure 1A), demonstrating that BRC-1 is not required for intersister/intrachromatid crossover or noncrossover repair. Notably, the overall proportion of crossover progeny among recombinants identified was increased at all timepoints scored (Figure 1A), suggesting that BRC-1 functions in *C. elegans* meiosis to repress intersister/intrachromatid crossover events.

**Figure 1.**
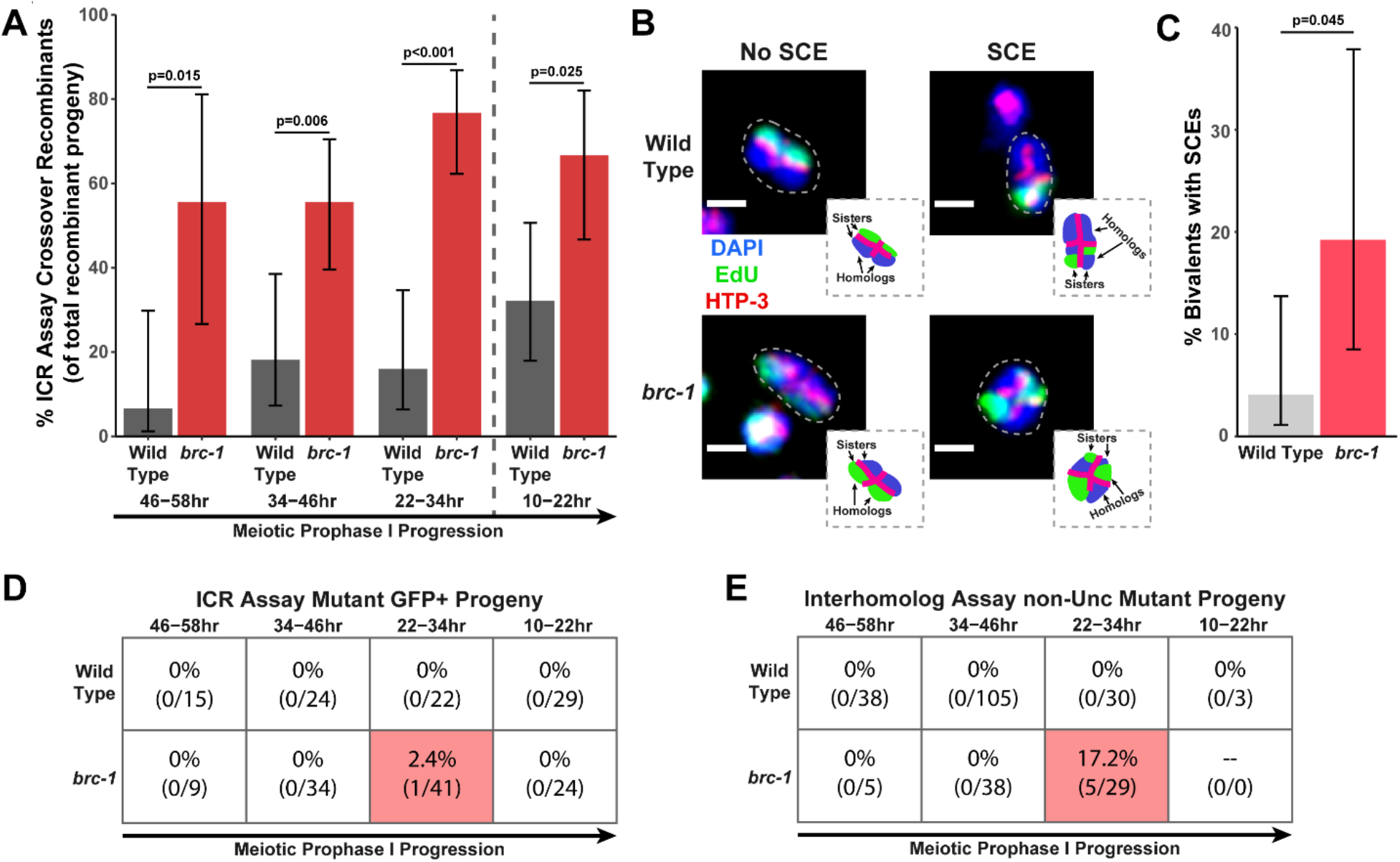
BRC-1 represses intersister crossovers and error-prone repair. A) Bar plot displaying the percent of crossover recombinant progeny identified in wild type and *brc-1* ICR assays out of all recombinant progeny scored. Frequencies of recombinants identified overall in ICR assays is displayed in Supplemental Figure 1A. B) Images of wild type and *brc-1(xoe4)* mutant bivalent chromosomes displaying an absence or presence of SCEs. Scale bars represent 1μm. Dashed bordered insets contain cartoon depictions of the SCE and non-SCE bivalents which are outlined with dashed lines in the images to aid in visualizing exchange events. C) Frequency of SCEs identified among wild type (n=49) or *brc-1* mutant (n=26) bivalents scored. D-E) Tables displaying the percent of sequenced GFP+ progeny in wild type and *brc-1* ICR assays (D) or non-Unc progeny IH assays (E) which showed signatures of mutagenic repair. Numbers in parentheses indicate the number of mutant worms out of the total number of sequenced progeny. Shaded boxes indicate timepoints in which mutant progeny were identified. The overall frequency of interhomolog assay non-Unc progeny is displayed in Supplemental Figure 2A-B. In all panels, error bars represent 95% Binomial confidence intervals, dashed vertical lines delineate between timepoints within the interhomolog window (22-58hr post heat shock) and non-interhomolog window (10-22hr post heat shock), and p values were calculated using Fisher’s Exact Test. **Figure 1 – source data 1. The source data for** Figure 1A, 1D **are provided.** [Figure 1 source data 1.xlsx]. The total number of ICR assay progeny with GFP+ or non-GFP+ phenotypes are listed. Wild type data I shared with Figure 2 and Supplemental Figure 1. **Figure 1 – source data 2. The source data for** Figure 1C **is provided.** [Figure 1 source data 2.xlsx]. The number of scorable chromatid pairs with SCE or no SCE events (no_SCE) are listed for each image assessed in generating this dataset. Wild type data is shared with Figure 2. **Figure 1 – source data 3. The source data for** Figure 1E **is provided.** [Figure 1 source data 3.xlsx]. The total number of IH assay progeny with recombinant or mutant nonUnc phenotypes or Unc nonrecombinant phenotypes are listed. Wild type data is shared with Figure 2 and Supplemental Figure 2.

To confirm that intersister crossovers are more frequent in a *brc-1* mutant, we employed a recently developed cytological assay which utilizes EdU incorporation to visualize sister chromatid exchanges (SCEs) in compacted diakinesis chromosomes (Figure 1B) (Almanzar *et al*. 2021, 2022). Notably, this cytological assay detects SCEs from endogenous SPO-11 induced DSBs. While SCEs are found in only 4.1% of bivalents in a wild type background (2/49 bivalents scored, 95% Binomial CI 1.1-13.7%) (Almanzar *et al*. 2021), we detected SCEs at an elevated rate of 19.2% in a *brc-1(xoe4)* mutant (Figure 1B-1C, 5/26 bivalents scored, 95% Binomial CI 8.5-37.9%, Fisher’s Exact Test p=0.045). When we compared the levels of SCEs cytologically identified with the frequency of ICR assay crossovers generated from Mos1-induced DSBs within the interhomolog window, the elevated frequency of SCEs (4.7 fold increase) closely mirrored the relative increase in crossovers as a proportion of all recombinants observed in the *brc-1* mutant ICR assay (4.6 fold increase). Taken together, these results demonstrate that BRC-1 functions to suppress intersister crossover recombination during *C. elegans* meiosis for both SPO-11-induced DSBs as well as Mos1-induced DSBs.

### BRC-1 is not required for interhomolog recombination

Since BRC-1 acts to suppress crossover recombination between sister chromatids, we next assessed if *brc-1* mutants exhibit defects in interhomolog recombination, including interhomolog crossovers. To assess the overall rates of interhomolog noncrossover and crossover recombination, we employed an established interhomolog (IH) recombination assay (Rosu *et al*. 2011) which enables: 1) controlled generation of a single DSB in meiotic nuclei via heat-shock inducible Mos1 excision (Robert and Bessereau 2007); 2) identification of interhomolog DSB repair of the induced DSB by reversion of an uncoordinated movement ‘Unc’ phenotype (non-Unc progeny, see Methods); and, 3) delineation of interhomolog noncrossover and crossover repair outcomes (see Methods). Notably, DSB repair in the IH assay which produces in-frame insertions or deletions can also yield non-Unc progeny which are phenotypically indistinguishable from noncrossover recombinants (Robert *et al*. 2008). While mutagenic repair in the IH assay is rare in a wild type context (Robert *et al*. 2008), *brc-1* mutants are known to incur small mutations more frequently (Kamp *et al*. 2020; Meier *et al*. 2021). We therefore sequenced the repaired *unc-5* locus of putative noncrossover non-Unc progeny in the IH assay to confirm whether the repaired sequence matched the homolog repair template or indicated mutations at the site of Mos1 excision (see Methods). Non-Unc progeny which we were unable to sequence were designated as ‘undetermined non-Unc’.

When we performed the IH assay in the *brc-1* mutant, we observed a significant increase in the proportion of non-Unc progeny only at the 22-34hr timepoint, which corresponds to the mid pachytene stage of meiosis and the end of the interhomolog window (Supplemental Figure 2A, Fisher’s Exact Test p<0.001). This result may indicate a slight delay in the rate of meiotic progression in *brc-1* mutants (Rosu *et al*. 2011). However, the overall frequency of non-Unc progeny was not elevated relative to wild type within the non-interhomolog window (Supplemental Figure 2A, 10-22hr post heat shock, Fisher’s Exact Test p=0.303), indicating that ablation of *brc-1* does not severely impact meiotic prophase I progression.

When we compared the ratio of crossover and noncrossover recombinant progeny within the interhomolog window between wild type and *brc-1* mutants, we saw that the frequency of interhomolog crossovers was not significantly altered (Supplemental Figure 2B, Fisher’s Exact Test p=0.515). This result mirrors recombination assays previously performed in *brc-1* mutants which provided no evidence for the presence of additional crossovers (Li *et al*. 2018). Thus, our data supports a role for BRC-1 in regulating crossover recombination specifically between sister chromatids.

### BRC-1 prevents mutagenic DNA repair during the mid-pachytene stage

In both the ICR and IH assays performed in *brc-1* mutants, we identified progeny which exhibited molecular signatures of mutagenic DSB repair at the Mos1 excision site (Figure 1D-E, Supplemental Figure 1A, Supplemental Figure 2A). These events were only identified within the 22-34hr timepoint, which is composed of nuclei in mid pachytene at the time of Mos1 excision. In the ICR assay, mutants were identified as 2.4% (95% Binomial CI 0.4-12.5%) of all sequenced GFP+ progeny at the 22-34hr time point. In the IH assay, 13.2% (95% Binomial CI 7.6-34.5%) of all sequenced non-Unc progeny at the 22-34hr time point were identified as mutant (Figure 1D-E). Notably, we only sequenced GFP+ and non-Unc progeny in the ICR and IH assays respectively. The frequency of error prone pathway utilization in a *brc-1* mutant is therefore likely much greater than our results suggest, as we could not detect mutations which disrupt the GFP or *unc-5* open reading frames.

Of the meiotic lesions we identified among *brc-1* IH assay progeny (see Methods), 75% (3/4 mutations) exhibited one or more complementary nucleotides on both ends of the deletion (Supplemental Figure 3B). Further, the single mutant identified among *brc-1* ICR assay GFP+ progeny displayed a particularly striking duplication joined at a position sharing microhomology (Supplemental Figure 3A). Regions of microhomology present on either end of small (<50bp) deletions and templated insertions are characteristic of theta mediated end joining (TMEJ) (Van Schendel *et al*. 2015). A previous study demonstrated that the rate of TMEJ-mediated germline mutagenesis is elevated in *brc-1* mutants (Kamp *et al*. 2020). Our data is therefore concordant with elevated TMEJ engagement in *brc-1* mutants and further reveals that the function of BRC-1 in preventing mutagenic repair events is specifically vital in the mid-pachytene stage of meiotic prophase I.

### SMC-5/6 restricts intersister crossovers

The SMC-5/6 DNA damage complex has been hypothesized to function in homolog-independent DSB repair in *C. elegans* (Bickel *et al*. 2010). To directly assess the functions of the structural maintenance of chromosomes 5/6 (SMC-5/6) complex in homolog-independent DSB repair, we performed the ICR assay in the *smc-5(ok2421)* null mutant. The *smc-5(ok2421)* deletion allele disrupts the final 6 exons of the 11 exons in the *smc-5* coding sequence and prevents SMC-5/6 complex assembly, as evidenced by both biochemical and cytological experiments (Bickel *et al*. 2010). SMC-5/6 is therefore not required for viability in *C. elegans*, unlike many other organisms (Aragón 2018). Similar to the *brc-1* mutant, we found that the frequency of GFP+ progeny in the ICR assay was elevated at all timepoints scored in *smc-5(ok2421)* null mutants (Supplemental Figure 1B). As mentioned above and in the Methods, this result does not necessarily represent an absolute increase in the rate of intersister/intrachromatid recombination (see Methods). Importantly, we did identify both crossover and noncrossover recombinants at all timepoints scored, demonstrating that SMC-5/6 is not required for noncrossover nor crossover homolog-independent repair (Supplemental Figure 1B).

To determine if SMC-5/6 regulates engagement of intersister/intrachromatid recombination outcomes, we examined the proportion of *smc-5* ICR assay crossover recombinants as a proportion of all recombinants identified. While the proportion of crossovers was not significantly different than wild-type within the individual 12-hour timepoints we scored (Figure 2A), the frequency of crossover recombinants in *smc-5* mutants was significantly elevated within the interhomolog window overall (Figure 2B, Fisher’s Exact Test p=0.037). Thus, our data suggests that a function of SMC-5/6 is to prevent homolog-independent crossovers arising from DSBs induced in early stages of meiotic prophase I. To cytologically affirm the results of our ICR assay, we assessed the frequency of SCEs in *smc-5(ok2421)* mutants by examining EdU labeled chromatids at diakinesis. Mutants for *smc-5* are known to have defects in chromosome compaction and produce misshapen bivalents (Bickel *et al*. 2010; Hong *et al*. 2016). These defects made the majority of bivalents uninterpretable in the EdU labeling assay. Nonetheless, even among a limited sample, we identified SCEs in 50% of scored bivalents (Figure 2C-D, 3/6 bivalents scored, 95% Binomial CI 18.8-81.2%, Fisher’s Exact Test p=0.007) as compared to only 4.1% of wild type bivalents (2/49 bivalents scored, 95% Binomial CI 1.1-13.7%) (Almanzar *et al*. 2021). This EdU labeling data in the *smc-5(ok2421)* null mutant represents a 12.2 fold increase in the rate of SCEs, which is notably more extreme than the 2.1 fold increase in the proportion of crossover recombinants observed in the IH window in our *smc-5* ICR assay data. Nevertheless, both our ICR assay and EdU labeling experiments support a function for SMC-5/6 in repressing intersister crossing over during *C. elegans* meiosis.

**Figure 2.**
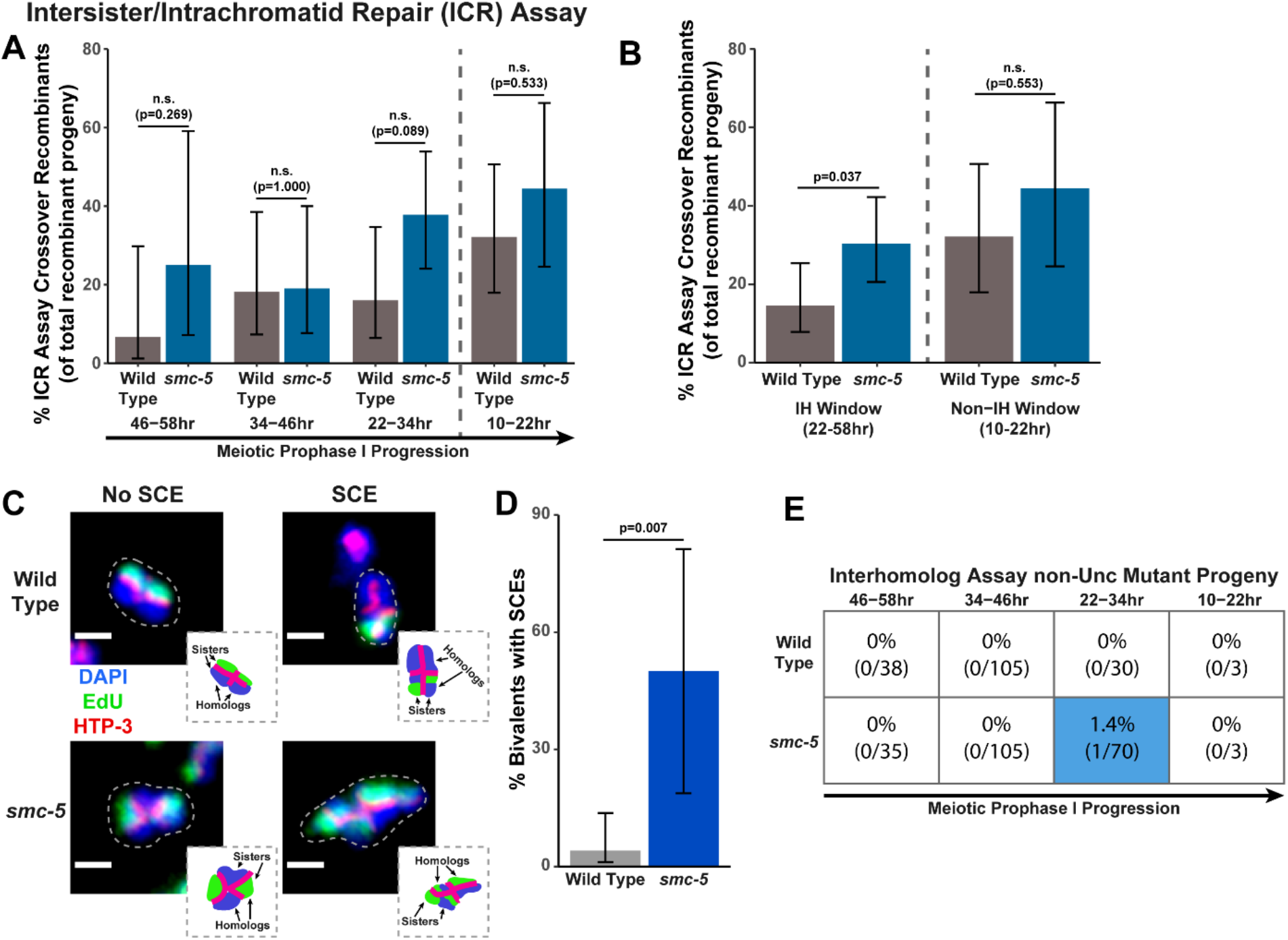
SMC-5/6 represses intersister crossovers. A) Bar plot displaying the percent of crossover recombinant progeny identified in wild type and *smc-5* ICR assays out of all recombinant progeny scored within individual 12 hour timepoint periods. Frequencies of recombinants identified overall in ICR assays is displayed in Supplemental Figure 1B. B) Bar plot displaying the percent of crossover recombinant progeny identified in wild type and *smc-5* ICR assays out of all recombinant progeny scored within the interhomolog window (22-58hr post heat shock) and non-interhomolog window (10-22hr post heat shock). C) Images of wild type and *smc-5(ok2421)* mutant bivalent chromosomes displaying an absence or presence of SCEs. Scale bars represent 1μm. Dashed bordered insets contain cartoon depictions of the SCE and non-SCE bivalents which are outlined with dashed lines in the images to aid in visualizing exchange events. D) Frequency of SCEs identified among wild type (n=49) or *smc-5(ok2421)* mutant (n=6) bivalents scored. E) Table displaying the percent of sequenced non-Unc progeny in wild type and *smc-5* IH assays which showed signatures of mutagenic repair. Numbers in parentheses indicate the number of mutant worms out of the total number of sequenced progeny. Colored boxes indicate timepoints in which mutant progeny were identified. The overall frequency of interhomolog assay non-Unc progeny is displayed in Supplemental Figure 2C-D. In all panels, error bars represent 95% Binomial confidence intervals, dashed vertical lines delineate between timepoints within the interhomolog window (22-58hr post heat shock) and non-interhomolog window (10-22hr post heat shock), and p values were calculated using Fisher’s Exact Test. **Figure 2 – source data 1. The source data for** Figure 2A **is provided.** [Figure 2 source data 1.xlsx]. The total number of ICR assay progeny with GFP+ or non-GFP+ phenotypes are listed. Wild type data is shared with Figure 1 and Supplemental Figure 1. **Figure 2 – source data 2. The source data for** Figure 2D **is provided.** [Figure 2 source data 2.xlsx]. The number of scorable chromatid pairs with SCE or no SCE events are listed for each image assessed in generating this dataset. Wild type data is shared with Figure 1. **Figure 2 – source data 3. The source data for** Figure 2E **is provided.** [Figure 2 source data 3.xlsx]. The total number of IH assay progeny with nonUnc phenotypes or Unc nonrecombinant phenotypes are listed. Wild type data is shared with Figure 1 and Supplemental Figure 2.

### SMC-5/6 is not required for interhomolog recombination

To determine if the SMC-5/6 complex regulates interhomolog recombination, we performed the IH assay in the *smc-5(ok2421)* null mutant. We identified both interhomolog crossover and noncrossover recombinants in the IH assay (Supplemental Figure 2C), indicating that SMC-5/6 is not required for either of these recombination pathways. Similar to *brc-1* mutants, we noted elevated non-Unc progeny at the 22-34hr time point in *smc-5* mutants, implying that meiotic prophase progression may be slightly delayed when SMC-5/6 function is lost (Supplemental Figure 2C, Fisher’s Exact Test p<0.001). Notably, non-Unc progeny were not increased in the non-interhomolog window in *smc-5* mutants, suggesting that the progression of meiotic prophase I was not drastically altered in this genetic context (Supplemental Figure 2C, Fisher’s Exact Test p=1.000). The proportion of crossover recombinants among all recombinants identified also was not altered in an *smc-5* mutant (Supplemental Figure 2D, Fisher’s Exact Test p=0.495). Thus, our data does not support a function for SMC-5/6 in ensuring efficient interhomolog recombination.

Among all sequenced ICR and IH assay GFP+ and non-Unc progeny isolated in *smc-5* mutants, we identified only one mutagenic DSB repair event at the 22-34hr timepoint of the IH assay (Figure 2E, Supplemental Figure 2C, Supplemental Figure 3B). Moreover, the frequency of *smc-5* non-Unc mutants which we detected at this timepoint (1.32% of all sequenced non-Unc progeny, 95% Binomial CI 0.2-7.1%) is lower than the frequency observed in *brc-1* mutants (Fisher’s Exact Test p=0.015). Previously, profiling of meiotic mutagenic DNA repair events in *smc-6* mutants revealed that large structural variations are a primary class of mutations which arise in SMC-5/6 deficient germlines (Volkova *et al*. 2020). In our *smc-5* ICR and IH assays, a greater frequency of DSBs may have been resolved by mutagenic repair, but if these products disrupted the coding sequence in GFP or *unc-5* respectively, then they would have escaped detection in our assays.

### BRC-1 promotes the formation of long homolog-independent noncrossover conversion tracts

Since we identified functions for BRC-1 and SMC-5/6 in regulating intersister crossover recombination, we wanted to determine if recombination intermediate processing is altered in *brc-1* and *smc-5* mutants. Evaluation of sequence conversions have informed much of our understanding of recombination intermediate processing (Szostak *et al*. 1983; Pâques and Haber 1999; Marsolier-Kergoat *et al*. 2018; Ahuja *et al*. 2021). The ICR assay was engineered to contain multiple polymorphisms spanning 12bp to 567bp 3’ from the site of Mos1 excision, enabling conversion tract analysis of homolog-independent recombination (Toraason *et al*. 2021a). In a wild type context, 74.2% of ICR assay noncrossover conversion tracts within the interhomolog window are ‘short’, which we define as tracts with a sequence conversion only at the most proximal polymorphism 12bp downstream from the site of Mos1 excision (Figure 4A, 4C, wild type 74.2% short tracts 95% CI 62.6-83.3%). In contrast to 74.2% of wild type noncrossover tracts during the interhomolog window being classified as ‘short’, 96.6% of *brc-1* noncrossover tracts during the interhomolog window were ‘short’ (*brc-1* interhomolog window 96.6% short tracts 95% CI 82.8-99.4%, Fisher’s Exact Test p=0.010). During the non-interhomolog window, a null mutation of *brc-1* had no effect on the proportion of ‘short’ noncrossover tracts (Figure 3A, 3C, wild type 72.7% short tracts 95% CI 51.8-86.8%; *brc-1* 87.5% short tracts 95% CI 52.9-97.8%, Fisher’s Exact Test p=0.638), thereby indicating that BRC-1 likely affects the mechanisms of noncrossover formation only during the interhomolog window.

**Figure 3.**
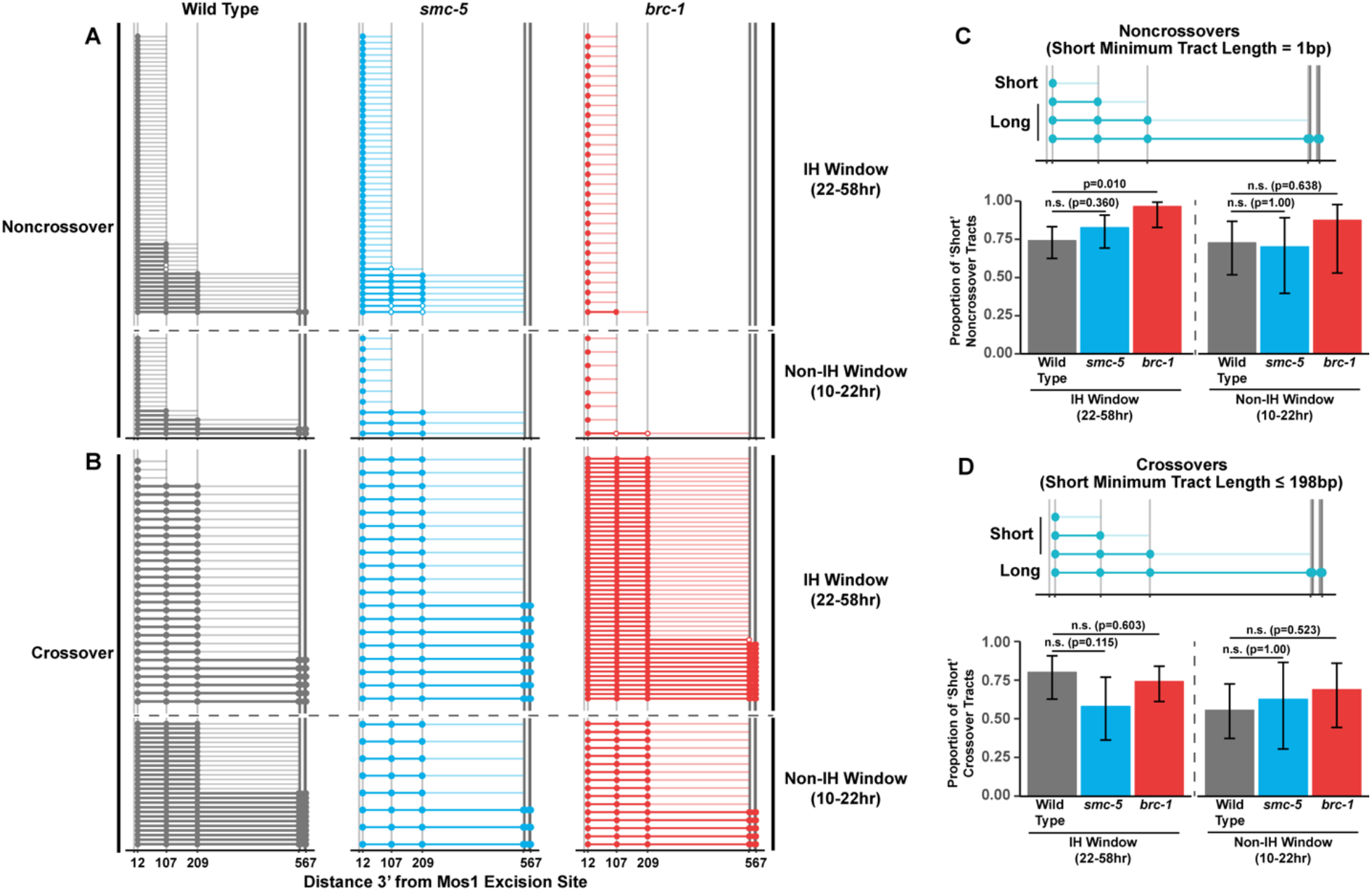
BRC-1 is required for long noncrossover gene conversion. A-B) Plots of conversion tracts sequenced from recombinant ICR assay loci. Vertical grey lines indicate the positions of polymorphisms in the ICR assay with bp measurements given 3’ relative to the site of Mos1 excision (Toraason *et al*. 2021a; b). Each horizontal line represents a single recombinant sequenced, ordered from smallest tract to largest tract within the interhomolog and non-interhomolog windows. Filled in points represent fully converted polymorphisms, while points with white interiors represent heteroduplex DNA sequences identified in conversion tracts. High opacity horizontal lines within plots represent the minimum conversion tract length, or the distance from the most proximal to the most distal converted polymorphisms. Low opacity horizontal lines indicate the maximum conversion tract, extending from the most distal converted polymorphism to its most proximal unconverted polymorphism. Tracts from noncrossover recombinants are displayed in A, while tracts from crossover recombinants are displayed in B. C-D) Frequency of short noncrossover tracts (C, minimum tract length 1bp converted at only the 12bp polymorphism) or short crossover tracts (D, minimum tract length 198bp) as a proportion of all tracts identified from progeny laid within the interhomolog and non-interhomolog windows. Error bars represent the 95% binomial confidence intervals of these proportions and p values were calculated using Fisher’s Exact Test. Diagrams above bar plots depict the sizes of tracts considered ‘long’ or ‘short’ in each respective group. In all panels, dashed grey lines delineate between the interhomolog window (22-58hr post heat shock) and non-interhomolog window (10-22hr post heat shock) timepoints. **Figure 3 – source data 1. The source data for** Figure 3 **is provided.** [Figure 3 source data 1.xlsx]. The polymorphism conversions scored in individual sequenced ICR assay conversion tracts are listed.

**Figure 4.**
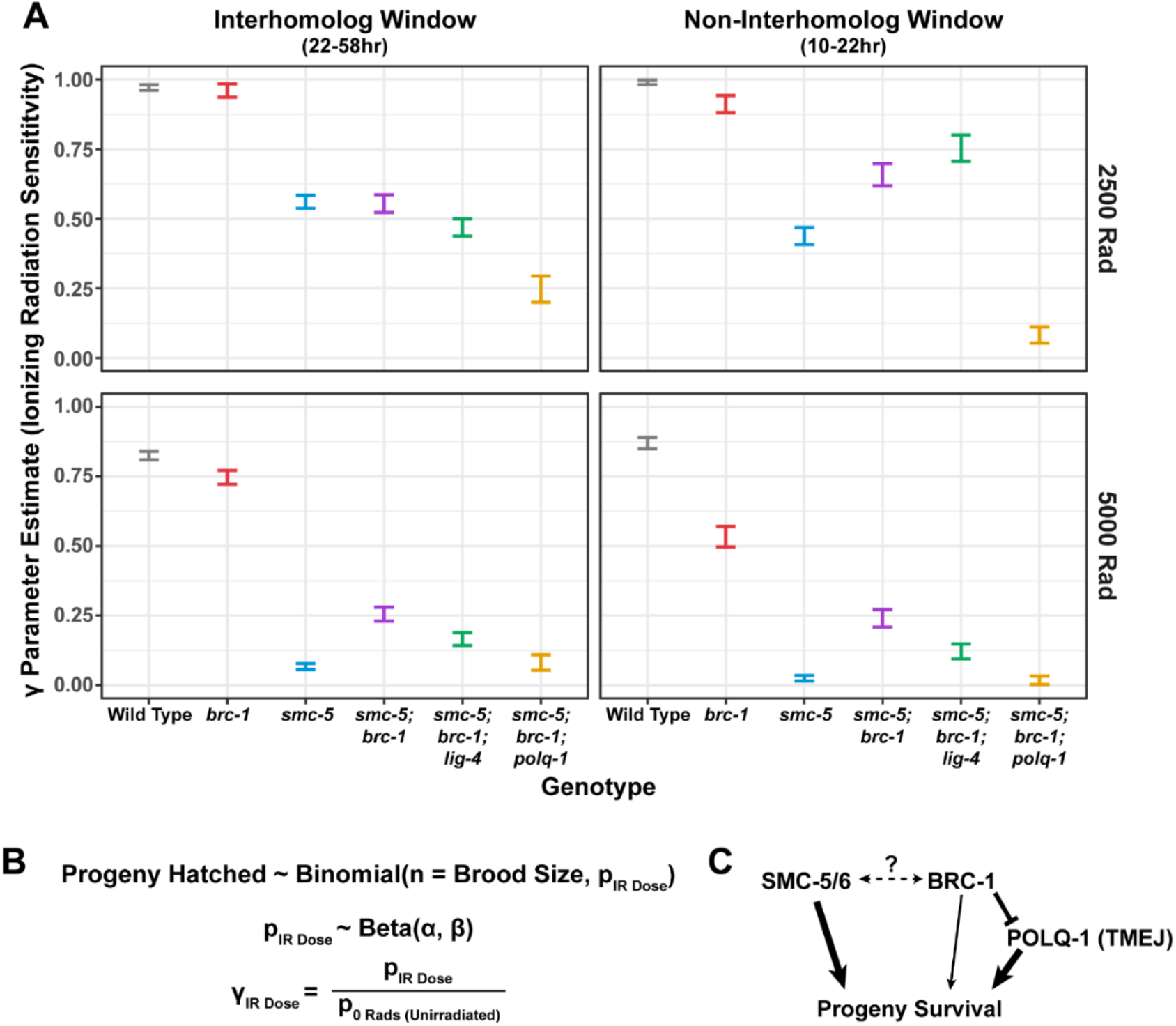
Interactions of SMC-5/6 and BRC-1 in meiotic DSB repair following irradiation. A) Gamma parameter estimates of genotype sensitivity to ionizing radiation of given doses. Vertical error bars represent the 95% credible interval of the gamma estimate for each genotype at the given dose of irradiation exposure. The brood viabilities of hermaphrodites used in this analysis are displayed in Supplemental Figure 4A and Supplemental Figure 4 source data 1. B) Outline of beta binomial model framework used to generate panel A. See Methods for details. C) Genetic interaction diagram inferred from estimates presented in panel A. SMC-5/6 and BRC-1 both contribute to progeny viability following meiotic exposure to exogenous DSBs. However, BRC-1 also inhibits error prone repair, which can compensate for the DSB defects of *smc-5* mutants when *brc-1* is also ablated. **Figure 4 source code 1. The source code for** Figure 4a **is provided.** [Figure 4 source code 1.R]. The R code utilized in performing the hierarchical statistical modeling in Figure 4a is provided. Code generating the posterior simulations displayed in Supplemental Figure 4B is also provided in this R script. **Figure 4 source code 2. The Rstan model fit for** Figure 4a **is provided.** [Figure 4 source code 2.rds]. The RStan output from the code in Figure 4 source data 1 is provided. **Figure 4 source data 1. The parameter estimates for** Figure 4a **are provided.** [Figure 4 source data 1.xlsx] Summary statistics of the posterior probability distributions for the p, alpha, and beta parameters and the gamma metric are listed. Statistics included are the mean, standard error of the mean (se_mean), standard deviation (sd), credible interval boundaries (2.5%, 25%,50%,75%,97.5%), effective sample size (n_eff), and the MCMC chain equilibrium metric *R̂*(Rhat).

We previously showed that wild type intersister/intrachromatid crossover conversion tracts in *C. elegans* tend to be larger than noncrossovers, with a median minimum conversion tract length (the distance from the most proximal to the most distal converted polymorphisms in bp) for intersister/intrachromatid crossovers being 198bp (Figure 3B) (Toraason *et al*. 2021a). Based on this median length for intersister/intrachromatid crossovers, we defined ‘short’ ICR assay crossover tracts as ≤198bp in length. We found that the proportion of ‘short’ crossover tracts was not altered by *brc-1* mutation within the interhomolog window (Figure 3B, 3D, wild type 80.0% short tracts 95% CI 62.7-90.5%; *brc-1* 74.1% short tracts 95% CI 61.1-83.9%, Fisher’s Exact Test p=1.00) nor within the non-interhomolog window (Figure 3B, 3D, wild type 55.6% short tracts 95% CI 37.3-72.4%; *brc-1* 68.8% short tracts 95% CI 44.4-58.8%, Fisher’s Exact Test p=0.657). Taken together, these results support a model in which BRC-1 regulates mechanisms of intersister/intrachromatid noncrossover recombination (and not crossover recombination) in the early stages of meiotic prophase I.

### SMC-5/6 does not regulate the extent of homolog-independent gene conversion

To assess if SMC-5/6 influences recombination intermediates, we compared *smc-5* mutant ICR assay conversion tracts to their wild type counterparts. We found that ICR assay noncrossover conversion tracts in *smc-5* mutants exhibited a similar proportion of ‘short’ tracts to wild type in both the interhomolog (Figure 3A, 3C, wild type 74.2% short tracts 95% CI 62.6-83.3%; *smc-5* 82.6% short tracts 95% CI 69.3-90.9%, Fisher’s Exact Test p=0.360) and non-interhomolog windows (Figure 3A, 3C, wild type 72.7% short tracts 95% CI 51.8-86.8%; *smc-5* 70% short tracts 95% CI 39.7-89.2%). Thus, SMC-5/6 does not have a strong effect on the extent of noncrossover gene conversion in intersister/intrachromatid repair.

When we compared the proportion of ‘short’ *smc-5* ICR assay crossover tracts to wild type, we similarly observed that there is no significant difference in the proportion of short and long crossover tracts in either the interhomolog (Figure 3B, 3D, wild type 80.0% short tracts 95% CI 62.7-90.5%; *smc-5* 57.9% short tracts 95% CI 36.3—76.9%, Fisher’s Exact Test p=1.00) or non-interhomolog windows (Figure 3B, 3D, wild type 55.6% short tracts 95% CI 37.3-72.4%; *smc-5* 62.5% short tracts 95% CI 30.6-86.3%, Fisher’s Exact Test p=1.00). Taken together, these results do not support a function for SMC-5/6 in regulating the extent of noncrossover and crossover gene conversion which yields functional GFP repair products.

In our wild type, *brc-1*, and *smc-5* ICR assay conversion tracts, we additionally noted multiple instances of heteroduplex DNA in our sequencing (Figure 3A, 3B). DNA heteroduplex is a normal intermediate when recombination occurs between polymorphic templates but is usually resolved by the mismatch repair machinery. Our observation of these events across genotypes suggests that at a low frequency, mismatch repair may fail to resolve heteroduplex DNA during the course *C. elegans* meiotic DSB repair.

### BRC-1 and SMC-5/6 genetically interact in resolving exogenous DSBs

To determine whether the regulation of homolog-independent DSB repair involves interactions between SMC-5/6 and BRC-1, we assessed how *smc-5(ok2421);brc-1(xoe4)* double mutants respond to DSBs. Since genetically balanced *smc-5;brc-1* double mutants can still acquire mutations and become progressively sterile over the course of a few generations and the ICR assay requires multiple cross steps (see Methods), we assessed the resilience of *smc-5*, *brc-1*, and *smc-5;brc-1* mutant gametes to exogenous DSBs induced by ionizing radiation to minimize the impact of this reproductive dysfunction phenotype. Accordingly, we treated wild type, *smc-5*, *brc-1*, and *smc-5;brc-1* mutant adult hermaphrodites with 0, 2500, or 5000 Rads of ionizing radiation, which induces DSBs, and assayed the resultant progeny derived from their irradiated oocytes for larval viability (Supplemental Figure 4A).

Importantly, we scored brood viability over a similar reverse time course as was done in our ICR and IH assays following irradiation (see Methods), enabling us to identify meiosis-stage specific DNA repair defects in these mutants. We noted variation in the brood viabilities of individual genotypes and individual hermaphrodites within genotypes (Supplemental Figure 4A), which indicates differences in baseline fertility even in unirradiated conditions. These baseline disparities posed a challenge in interpreting the effects of ionizing radiation on brood viability, as the resilience of an irradiated cohort will be affected by both underlying fertility defects as well as the effects of the exogenous DNA damage that we sought to quantify. To estimate the effect of ionizing radiation on brood viability and to account for inter-hermaphrodite variance in our analysis, we employed a hierarchical statistical modeling approach using our dataset (Figure 4B, see Methods). From this analysis, we calculated a metric termed ‘gamma’ for each genotype, representing the sensitivity of a given genotype to ionizing radiation (Figure 4B, see Methods). A gamma estimate of 1 indicates that irradiation has no effect on brood viability, while a gamma estimate of 0 indicates that all progeny of a genotype are inviable following irradiation.

To assess the differential sensitivities of *smc-5*, *brc-1*, and *smc-5;brc-1* mutants across meiotic prophase I, we compared the 95% credible intervals of the gamma estimates for each genotype within the interhomolog and the non-interhomolog windows for both moderate (2500 Rads) and high (5000 Rads) irradiation doses (Figure 4A). Across all irradiation doses and timepoints, we note that loss of *smc-5* conveys a greater sensitivity to exogenous DNA damage than loss of *brc-1* (Figure 4A), emphasizing that the SMC-5/6 complex prevents catastrophic defects following exogenous DNA damage induction. Moreover, the sensitivity of both single mutants to ionizing radiation is greater in the non-interhomolog window than in the interhomolog window (Figure 4A). This result demonstrates that meiotic cells are more dependent upon these complexes to resolve DSBs when the homolog is unavailable as a repair template.

At 2500 Rad of ionizing radiation, we found that mutation of both *smc-5* and *brc-1* differentially impacted radiation resilience within the interhomolog and non-interhomolog windows. In the interhomolog window, the *smc-5;brc-1* double mutant and *smc-5* single mutant gamma estimates overlap, indicating that loss of BRC-1 does not alter *smc-*5 mutant sensitivity at this timepoint (Figure 4A). Further, *brc-1* mutant gamma estimates are indistinguishable from wild type within the interhomolog window (Figure 4A); therefore, the absence of an interaction may reflect the dispensability of BRC-1 in early prophase I for progeny survival when DNA damage levels are not extreme. In the non-interhomolog window, however, we observe a striking resilience to exogenous DSBs in *smc-5;brc-1* double mutants as compared to *smc-5* single mutants (Figure 4A). This synthetic resilience is recapitulated across meiotic prophase I at 5000 Rads of ionizing radiation in *smc-5;brc-1* double mutants (Figure 4A). Thus, our data indicates that DNA damage sensitivity observed in *smc-5* mutants is enhanced by BRC-1-mediated functions.

BRC-1 is known to repress both TMEJ and NHEJ in multiple organisms, including *C. elegans* (Huen *et al*. 2010; Li *et al*. 2020; Kamp *et al*. 2020). We hypothesized that error prone repair pathways may be activated in *smc-5;brc-1* double mutants to resolve DSBs and abrogate the DNA repair defects associated with *smc-5* mutation. To test whether TMEJ and/or NHEJ contribute to the ionizing radiation resilience observed in *smc-5;brc-1* double mutants, we created *smc-5;brc-1;polq-1* and *smc-5;brc-1;lig-4* triple mutants which are defective in TMEJ and NHEJ respectively. We observed a striking effect at all radiation doses and timepoints scored in a *smc-5;brc-1;polq-1* mutant as compared to the *smc-5;brc-1* mutant. Even at the moderate dose of 2500 Rads, loss of POLQ-1 caused dramatic sensitization of *smc-5;brc-1* mutants to ionizing radiation (Figure 4A). This effect was particularly strong in the non-interhomolog window, where *smc-5;brc-1;polq-1* mutants were nearly sterile following ionizing radiation treatment regardless of irradiation dose (Figure 4A). Previous irradiation studies have shown that neither *polq-1* nor *brc-1;polq-1* mutation confer as severe of a radiation sensitivity phenotype as we observe in the *smc-5;brc-1;polq-1* triple mutant (Bae *et al*. 2020; Kamp *et al*. 2020). These results strongly indicate that *smc-5;brc-1* deficient germ cells exposed to exogenous DNA damage are dependent upon TMEJ for fertility.

In contrast to the dramatic effects on DSB repair produced in our *smc-5;brc-1;polq-1* mutant, we found that *smc-5;brc-1;lig-4* mutants exhibited only mild effects on radiation sensitivity compared to the *smc-5;brc-1* double mutant alone (Figure 4A). As loss of *lig-4* did not fully suppress the synthetic radiation resilience of *smc-5;brc-1* mutants, our experiments suggest that NHEJ is not a primary mechanism of DNA repair in meiotic nuclei when both SMC-5/6 and BRC-1 are lost. Taken together, the results of our irradiation analysis indicate that both SMC-5/6 and BRC-1 contribute to gamete viability following ionizing radiation treatment, with loss of SMC-5/6 having far greater consequences for the gamete than loss of BRC-1 (Figure 4C). As *brc-1* mutation confers synthetic resilience to radiation in *smc-5* mutants, we provide evidence that some functions of BRC-1 contribute to the meiotic DSB repair defects associated with *smc-5* mutation (Figure 4C). Further, we find that TMEJ is vital to radiation resilience in *smc-5;brc-1* mutants, suggesting that this pathway compensates for the DNA repair deficiencies incurred when SMC-5/6 and BRC-1 are both lost (Figure 4C). Repression of TMEJ by BRC-1 may therefore be deleterious to reproductive success in *smc-5* mutants by enabling more severe DNA repair errors to occur.

### BRC-1 localization is independent of SMC-5/6

To determine whether there is a co-dependency between BRC-1 and SMC-5/6 for localization, we first examined GFP::BRC-1 by immunofluorescence in both wild type and *smc-5* mutant germlines. Similar to previous studies (Janisiw *et al*. 2018; Li *et al*. 2018), we observed that BRC-1 localizes as a nuclear haze in the premeiotic tip through early pachytene and becomes associated with the synaptonemal complex during the progression of pachytene in wild type germlines (Supplemental Figure 5). In late pachytene, BRC-1 relocates to the short arms of the bivalents, where it can be visualized at diplotene as short tracks on the compacted chromosome arms (Supplemental Figure 5). When we examined *smc-5* mutants, the general pattern of GFP::BRC-1 localization across meiotic prophase was similar to wild type, except in the premeiotic tip where GFP::BRC-1 formed bright foci (Supplemental Figure 5). Given that BRD-1, the obligate heterodimeric partner of BRC-1, was found to form a similar localization in *smc-5* mutants (Wolters *et al*. 2014), the bright GFP::BRC-1 foci in the pre-meiotic tip likely mark BRC-1 localization to collapsed replication forks (Bickel *et al*. 2010; Wolters *et al*. 2014). Our data therefore indicate normal localization of BRC-1 does not require SMC-5/6.

To assess if BRC-1 changes localization in response to exogenous DSBs, we exposed wild type and *smc-5* mutant germlines to 5000 Rads of ionizing radiation and again examined germline GFP::BRC-1 by immunofluorescence. We found that the general pattern of GFP::BRC-1 localization appeared normal in both wild type and *smc-5* mutants following irradiation (Supplemental Figure 5). Taken together, our results suggest that BRC-1 localization is not altered following the induction of exogenous DSBs even when SMC-5/6 complex function is lost.

### SMC-5/6 localization is independent of BRC-1

To determine whether SMC-5/6 localization is dependent upon BRC-1, we generated an endogenous *smc-5* allele which codes for the auxin-inducible degron (AID*) and 3xFLAG epitope tags on the C terminus (*smc-5(syb3065[AID*::3xFLAG])*). The *smc-5(syb3065)* allele did not confer sensitivity to ionizing radiation nor an alteration in RAD-51 loading, suggesting that the tag does not impair SMC-5/6 complex function (Supplemental Figures 6A and 7). We examined the localization of SMC-5::AID*::3xFLAG in both wild type and *brc-1* mutants (Supplemental Figure 6B). We observed that, similar to a prior study (Bickel *et al*. 2010), SMC-5/6 is present in meiotic nuclei throughout prophase I. Notably, we found that SMC-5 staining in early and mid-pachytene was primarily localized to the chromosome axis, marked with HTP-3 (Supplemental Figure 6B; see Methods). This localization pattern was altered in the transition to diplotene, when we observed that SMC-5 localizes to the chromatin on the compacting bivalent chromosomes, matching previous analyses (Supplementary Figure 6B) (Bickel *et al*. 2010). The pattern of SMC-5 localization was not disrupted in a *brc-1* mutant, and similarly was not altered following exposure to 5000 Rads of ionizing radiation (Supplementary Figure 6B). Thus, the localization of SMC-5/6 does not depend upon the activity of BRC-1 and is not altered following induction of exogenous DNA damage at the levels we tested.

## Discussion

Meiotic cells must coordinate DNA repair pathway engagement to ensure both formation of interhomolog crossovers and repair of all DSBs. The highly conserved proteins SMC-5/6 and BRC-1 promote accurate DSB repair, but the specific DNA repair outcomes that these proteins regulate have remained unclear. We find that SMC-5/6 and BRC-1 both act to repress intersister crossovers, and further demonstrate that BRC-1 specifically influences noncrossover intermediate processing. We also observe that mutants for *brc-1* incur DNA repair defects at mid pachytene, as evidenced by increased engagement of error prone repair pathways. By comparing the germ cell resilience of *smc-5*, *brc-1,* and *smc-5;brc-1* mutants to ionizing radiation, we show that SMC-5/6 and BRC-1 are especially important for DSB repair in late meiotic prophase I. Further, we reveal that BRC-1 enhances the meiotic DNA repair defects of *smc-5* mutants and provide evidence that this interaction is in part underpinned by BRC-1 dependent repression of TMEJ. Taken together, our study illuminates specific functions and interactions of highly conserved DNA repair complexes in promoting germline genome integrity.

### Functions of BRC-1 in *C. elegans* meiotic DNA repair

The work presented in this study demonstrates that meiotic cells deficient in BRC-1 exhibit multiple DNA repair defects, including reduced noncrossover conversion tract length, elevated rates of intersister crossovers, and engagement of error prone DSB repair mechanisms at the mid-pachytene stage. What functions of BRC-1 may underpin these phenotypes? Accumulating evidence in other model systems supports roles for BRCA1 in regulating many early steps in recombination including DSB resection, strand invasion, and D-loop formation (Chen *et al*. 2008; Chandramouly *et al*. 2013; Cruz-García *et al*. 2014; Zhao *et al*. 2017; Kamp *et al*. 2020). We propose that perhaps some of these functions of BRC-1 are conserved in *C. elegans*.

While a growing body of research in budding yeast, mammalian systems, and *Arabidopsis* suggests that SDSA is the primary pathway for the formation of noncrossovers in meiosis (Hunter 2015; Marsolier-Kergoat *et al*. 2018; Ahuja *et al*. 2021) and that processing of joint molecular intermediates can generate noncrossovers during Drosophila meiosis (Crown *et al*. 2014), the mechanisms by which *C. elegans* noncrossover recombination occurs is unknown. Using the ICR assay, we find that the *brc-1* mutation affects the extent of ICR assay noncrossover gene conversion, but not crossover gene conversion, suggesting that homolog-independent noncrossovers arise from a distinct intermediate or undergo differential processing from crossovers in *C. elegans*. This result is consistent with a model in which either SDSA or joint molecule dissolution is a primary mechanism of intersister noncrossover recombination in the *C. elegans* germline (Figure 5).

**Figure 5.**
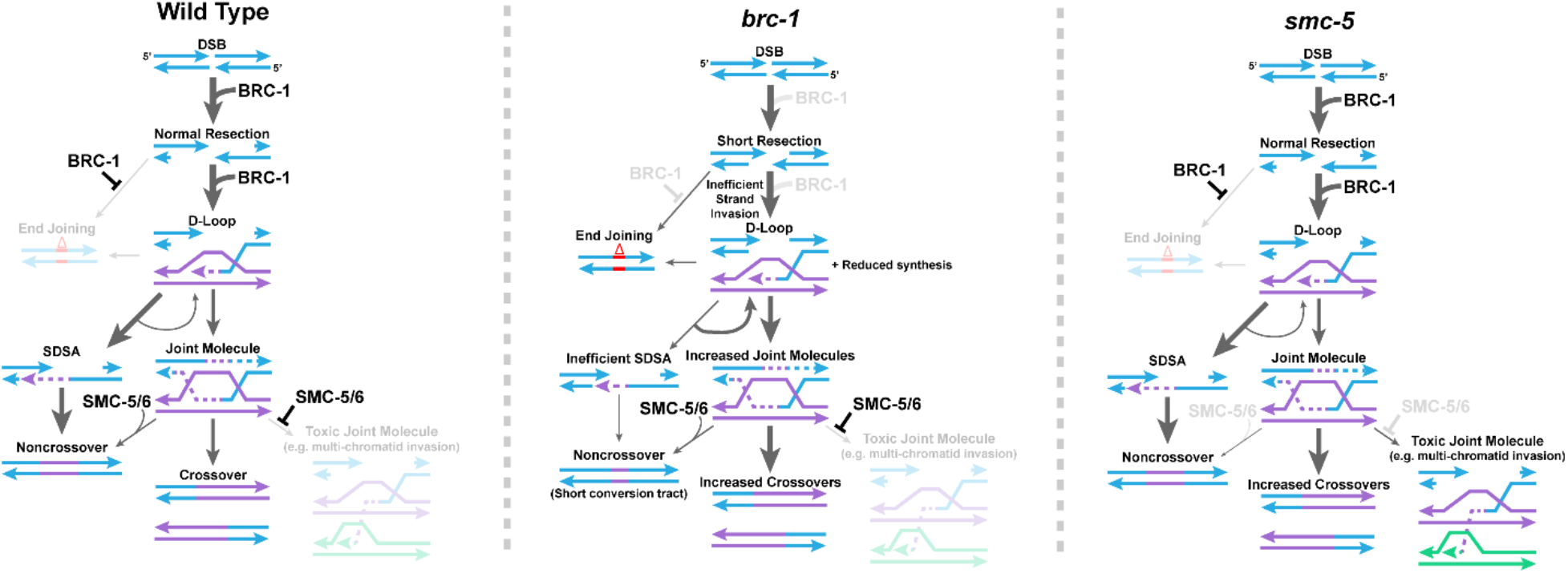
Model of BRC-1 and SMC-5/6 function in *C. elegans* intersister DSB repair. Displayed is a proposed model for the functions of BRC-1 and SMC-5/6 in regulating intersister DSB repair in the *C. elegans* germline. Under wild type conditions, BRC-1 promotes efficient resection of the damaged chromatid (blue) and facilitates strand invasion and extension with the sister chromatid (purple). BRC-1 also inhibits TMEJ either through direct antagonism of this pathway or indirectly by promoting efficient recombination. Following strand extension, the majority of D-loop intermediates are dissolved and repaired through SDSA, which is efficient due to BRC-1 promoted resection of the second end of the DSB. A minority of D-loops will proceed to form joint molecules, which may potentially be preferentially resolved as noncrossovers via the action of SMC-5/6 or as crossovers in an SMC-5/6 independent manner. In addition, SMC-5/6 inhibits the formation of toxic joint molecule intermediates, such as multi-chromatid joint molecules. In a *brc-1* mutant, DSBs are not resected to wild type levels and strand invasion is inefficient. Reduced resection limits the efficiency of second end capture in SDSA, reducing noncrossovers through this pathway. Further, limited strand extension reduces the extent of gene conversion in noncrossovers generated by successful SDSA or joint molecule dissolution. Failure in SDSA leads to increased DSB reinvasion of repair templates, contributing to the tandem duplications observed in mutants for BRCA1 (Chandramouly *et al*. 2013; Kamp *et al*. 2020). In addition, either due to absence of direct inhibition by BRC-1 or inefficiencies in recombination, end joining (particularly TMEJ) becomes activated to resolve DSBs. However, reduced resection does not inhibit joint molecule formation, leading to more of these intermediates which are preferentially resolved as crossovers. Finally, in an *smc-5* mutant, early steps in DSB repair proceed normally. However, absence of SMC-5/6 results in unconstrained joint molecule formation, including toxic intermediates. Failure in SMC-5/6 action to promote noncrossover repair further increases the proportion of joint molecules which are resolved as crossovers.

The size of an SDSA or dissolution of a noncrossover conversion tract depends upon the extent of heteroduplex DNA present following strand annealing, which is primarily determined by the length of DNA strand extension (Keelagher *et al*. 2011; Guo *et al*. 2017; Marsolier-Kergoat *et al*. 2018). Human BRCA1 promotes strand invasion and D-loop formation (Zhao *et al*. 2017), which may influence the efficiency of strand extension. Our conversion tract data raises the possibility that BRC-1 influences the formation and/or stability of strand invasion intermediates, thereby promoting the formation of long ICR assay noncrossover gene conversion events (Figure 5).

Our data also demonstrate that *brc-1* mutants exhibit elevated intersister crossovers. If BRC-1 only functions to promote strand invasion and D-loop formation, then we expected *brc-1* mutation to reduce intersister crossovers and not increase their occurrence. Previous studies have also suggested that BRCA1/BRC-1 regulates DSB resection, and we propose that this function better accounts for the observed increase in intersister crossovers (Chen *et al*. 2008; Cruz-García *et al*. 2014). Specifically, studies have posited that BRCA1-promoted long range DSB resection may be important for the efficiency of SDSA by ensuring sufficient single stranded DNA is exposed on the second end of the DSB to facilitate strand annealing (Chandramouly *et al*. 2013; Kamp *et al*. 2020). While sufficient resection may be critical in resolving noncrossovers, work in budding yeast has shown that long range resection is not required for the efficient formation of joint molecules (Zakharyevich *et al*. 2010). Thus, reduced length of DNA resection due to a *brc-1* mutation may impede SDSA and therefore increase the probability that DSBs will form joint molecule intermediates, thereby promoting intersister crossover outcomes.

Reduced resection in conjunction with inefficient strand invasion and synthesis during recombination may further explain the ectopic engagement of TMEJ observed in *brc-1* mutants (Kamp *et al*. 2020). Short range resection provides sufficient substrate for TMEJ (Ramsden *et al*. 2022), which in combination with inefficient homology search may provide more opportunity for TMEJ engagement. BRC-1 is also required in late meiotic prophase I for the loading and/or maintenance of RAD-51 at irradiation induced DSBs (Janisiw *et al*. 2018; Li *et al*. 2018). Defects in RAD-51 localization may further exacerbate the likelihood of error prone DSB repair at these meiotic stages. Overall, our data is consistent with a model in which BRC-1 promotes multiple DSB repair steps, including resection and the formation of early strand invasion intermediates, to facilitate intersister noncrossover repair (Figure 5).

### Functions of SMC-5/6 in *C. elegans* meiotic DSB repair

Our experiments demonstrate that SMC-5/6 acts to repress intersister crossover recombination in the early stages of meiotic prophase I. We do not find evidence, however, of prominent roles for SMC-5/6 in regulating ICR assay conversion tracts nor limiting error prone repair outcomes. These relatively subtle phenotypes appear at first incongruous with the known severe defects associated with loss of SMC-5/6 in *C. elegans*, which include chromosome fragmentation, large mutations, and transgenerational sterility (Bickel *et al*. 2010; Volkova *et al*. 2020). The ICR and IH assay experiments, however, are limited to the detection of DSB repair outcomes which encode a functional protein product. Thus, many of the severe mutations associated with SMC-5/6 deficiency may disrupt the coding sequence in the ICR or IH assays and therefore escape our detection (Volkova *et al*. 2020).

In budding yeast, Smc5/6 prevents the accumulation of toxic interchromosomal attachments and recombination intermediates (Chen *et al*. 2009; Xaver *et al*. 2013; Lilienthal *et al*. 2013; Copsey *et al*. 2013; Bonner *et al*. 2016; Peng *et al*. 2018) Prior evidence in *C. elegans* suggests that some of these functions are likely conserved, as double mutants for *smc-5* and the BLM helicase homolog *him-6* are sterile and display chromatin bridges indicative of persistent interchromosomal attachments (Hong *et al*. 2016). This synthetic phenotype suggests that these two complexes act in parallel to prevent the accumulation of joint molecules. A previous study (Almanzar *et al*. 2021) and the data we present here reveal that both SMC-5/6 and HIM-6 repress intersister crossovers. The synthetic sterility associated with loss of both SMC-5/6 and HIM-6 then may be a product of parallel functions for these proteins in limiting and/or resolving joint molecules. Although BLM is known to play multiple roles in regulating recombination, a core function of this helicase is in antagonism of joint molecule formation and promotion of noncrossover recombination (McVey *et al*. 2004; Weinert and Rio 2007; Schvarzstein *et al*. 2014). SMC-5/6 in *C. elegans* meiosis may therefore act as a second line of defense to ensure the elimination of inappropriate joint molecule intermediates which have formed more stable configurations (Figure 5). Under this model, we would expect accumulation of intersister joint molecules in an *smc-5* mutant and therefore elevated intersister crossovers, as observed in our *smc-5* ICR assay and EdU labeling experiments. Our observation that *smc-5* mutation does not alter ICR assay conversion tracts is also consistent with a model in which SMC-5/6 influences recombination following joint molecule formation. While the specific mechanisms by which SMC-5/6 may influence recombination intermediate formation or resolution remain unclear, recent work has shown that SMC-5/6 is capable of DNA loop-extrusion, indicating a function by which the complex may organize chromatin to facilitate efficient DSB repair (Pradhan *et al*. 2022). Specific subunits of SMC-5/6 also exhibit enzymatic function, such as the E3 SUMO ligase Nse2/Mms21 (Andrews *et al*. 2005), suggesting that SMC-5/6 may act to postranslationally modify target proteins to regulate DNA repair. Taken together, our data indicates that SMC-5/6 is not required for homolog-independent meiotic recombination and instead reveals a function for this complex in limiting crossover exchanges between sister chromatids.

### Temporal regulation of error-prone meiotic DSB repair

In both the ICR and IH assays we performed in *brc-1* mutants and in the IH assay we performed in *smc-5* mutants, we identified mutagenic repair events specifically at the 22-34hr timepoint, corresponding to oocytes in mid pachytene at the time of Mos1-excision induced DSB formation. Further, the repair events we identified frequently displayed microhomologies flanking the deletion site – a characteristic signature of TMEJ. While our dataset cannot definitively demonstrate that these events are the product of TMEJ, previous evidence and the nature of the break repair products strongly suggest that they originate from this pathway (Kamp *et al*. 2020). The limited temporal window in which we identified these events suggests that the activity of TMEJ may be relegated to later stages of meiotic prophase I. There are a number of important events which coincide at the mid/late pachytene transition of *C. elegans* meiosis, including a MAP kinase phosphorylation cascade, designation of interhomolog crossovers, a switch from RAD-50 dependence to independence for loading of RAD-51 to resected DNA, and loss of access to the homolog as a ready repair template (Church *et al*. 1995; Kritikou *et al*. 2006; Hayashi *et al*. 2007; Lee *et al*. 2007; Rosu *et al*. 2011; Yokoo *et al*. 2012; Nadarajan *et al*. 2016). These events may correspond to a switch in cellular “priorities” from ensuring interhomolog recombination to promoting repair of all residual DSBs even through error prone mechanisms. By repairing all residual DSBs (even in the wake of sequence errors), germ cells avoid catastrophic chromosome fragmentation during the meiotic divisions.

During the mid to late pachytene transition, an important function of BRC-1 (and to a lesser extent SMC-5/6) may be to prevent TMEJ either by antagonizing this pathway or facilitating efficient recombination. Our irradiation experiments revealed that both *brc-1* and *smc-5* mutant oocytes exhibit greater sensitivity to exogenous DNA damage in late stages of prophase I, suggesting that cellular requirements for efficacious DSB repair change during the transition to late pachytene. Moreover, during the late pachytene stage, several changes regarding BRC-1 occur: 1) BRC-1 protein localization changes; and, 2) BRC-1 is required to load (and/or stabilize) RAD-51 filaments (Janisiw *et al*. 2018; Li *et al*. 2018). We found that *brc-1* mutants incur mutations with characteristic TMEJ signatures specifically at the mid/late pachytene stage, suggesting that the changes in BRC-1 localization and function at this stage may coincide with changes in the availability and/or regulation of error prone repair mechanisms. Our irradiation experiments demonstrated that *smc-5;brc-1* double mutant oocytes throughout prophase I are dependent upon TMEJ DNA polymerase θ homolog *polq-1* for viability. If BRC-1 functions which repress TMEJ (Kamp *et al*. 2020) are specific to late prophase, then this result suggests that many DSBs in *smc-5;brc-1* mutants induced in early prophase may not be repaired until mid/late pachytene, when TMEJ is active. Spatiotemporal transcriptomic analysis has shown that *polq-1* is expressed throughout meiotic prophase I (Tzur *et al*. 2018). As we only identified error-prone resolution of DSBs induced at mid pachytene, our findings raise the possibility that BRC-1 independent mechanisms may repress TMEJ in early/mid pachytene. Our results in *brc-1* mutants therefore lay the groundwork for future research delineating the temporal regulation of error-prone meiotic DSB repair. Taken together, our study reveals that the engagement of error-prone and recombination DSB repair pathways are differentially regulated during the course of *C. elegans* meiotic prophase I.

### Interaction between BRC-1 and SMC-5/6 in meiotic DNA repair

Our irradiation experiments assessing the viability of *smc-5*, *brc-1*, and *smc-5;brc-1* mutant oocytes reveal that functional BRC-1 enhances the DNA repair defects of *smc-5* mutants. By further ablating error prone repair pathways, we also demonstrated that *smc-5;brc-1* mutants are dependent upon TMEJ for viability following irradiation. However, this genetic interaction does not coincide with changes in either SMC-5/6 or BRC-1 localization in respective mutants. Taken together, we suggest that the observed genetic relationships between BRC-1 and SMC-5/6 are likely not derived from direct physical interactions between these complexes, nor action on shared substrates, but rather arise from their respective sequential roles in regulation of DSB repair. A similar model was proposed by Hong *et al*. 2016 which postulated that early recombination defects in *brc-1* mutants may alleviate the toxic recombination intermediates formed in *smc-5;him-6* double mutants. We expand upon this model to demonstrate that this genetic relationship observed in *smc-5;him-6;brc-1* mutants is recapitulated in *smc-5;brc-1* double mutants, indicating that this interaction is not unique to the triple mutant context.

How might DNA repair defects in *brc-1* mutants ameliorate genomic instability associated with *smc-5* mutation? If *smc-5* mutants accumulate toxic joint molecules, then we would expect deficiencies in earlier recombination steps to limit the formation of these problematic intermediates and therefore alleviate the effects of *smc-5* mutation. Our analysis of homolog independent recombination in *brc-1* mutants revealed phenotypes which are consistent with this protein regulating both DSB resection and strand invasion. Work in budding yeast has shown that the additional ssDNA generated by long range resectioning of a DSB is used for homology search (Chung *et al*. 2010). Inefficient resection in *brc-1* mutants may reduce the extent of homology which could anneal to heterologous templates and contribute to toxic joint molecules (Figure 5). Conversely, resection defects of *brc-1* mutants could increase the risk for toxic recombination intermediates in *smc-5* mutants by limiting the efficiency of SDSA and therefore biasing DSBs to form joint molecules. However, compromised strand invasion and D-loop formation in *brc-1* mutants could also limit the capacity for DSBs to form multi-chromatid engagements. Finally, increased TMEJ activity in *smc-5;brc-1* mutants could resolve DSBs before they form recombination intermediates, thereby bypassing requirements for SMC-5/6 in DSB repair. In summation, our study reveals an interplay between BRC-1 and SMC-5/6 in regulating meiotic DSB repair.

## Acknowledgements

We thank the CGC (funded by National Institutes of Health P40 OD010440) for providing strains. We also thank J. Engebrecht for generously sharing strains carrying the *brc-1(xoe4)* and *GFP::brc-1* alleles. We thank C. Cahoon, A. Naftaly, and N. Kurhanewicz for thoughtful comments on this manuscript. We also thank G. Csankovszki for sharing antibodies for SMC-5 and SMC-6. This work was supported by the National Institutes of Health T32GM007413 and Advancing Science in America (ARCS) Foundation Award to E.T; National Institutes of Health R25HD070817 to A.S.; Genetics Training Grant 5T32M007464-42 to D.E.A.; a Pilot Project Award from the American Cancer Society, R35GM128804 grant from NIGMS, and start-up funds from the University of Utah to O.R.; and National Institutes of Health R00HD076165 and R35GM128890 to D.E.L. D.E.L. is also a recipient of a March of Dimes Basil O’Connor Starter Scholar award and Searle Scholar Award.

## Materials and Methods

### *Caenorhabditis elegans* strains and maintenance

*Caenorhabditis elegans* strains were maintained at 15°C or 20°C on nematode growth medium (NGM) plates seeded with OP50 bacteria. All experiments were performed in the N2 genetic background of *C. elegans* and animals were maintained at 20°C for at least two generations preceding an experiment.

Strains used in this study include:

N2 (wild type)

AV554 (*dpy-13(e184sd) unc-5(ox171::Mos1)*/ nT1 (qIs51) *IV*; KrIs14 (*phsp-16.48::MosTransposase*; *lin-15B*; *punc-122::GFP*) / nT1 (qIs51) *V*)

CB791 (*unc-5(e791) IV*),

DLW14 (*unc-5(lib1*[ICR assay *pmyo-3::GFP(-)*; *unc-119(+)*; *pmyo-2::GFP(Mos1)*]) *IV*; KrIs14 (*phsp-16.48::MosTransposase*; *lin-15B*; *punc-122::GFP*) *V*)

DLW23 (*smc-5(ok2421)*/mIn1 [*dpy-10(e128)* mIs14] *II*; *unc-5(lib1*[ICR assay *pmyo-3::GFP(-)*; *unc-119(+)*; *pmyo-2::GFP(Mos1)*]) *IV*; KrIs14 (*phsp-16.48::MosTransposase*; *lin-15B*; *punc-122::GFP*) *V*)

DLW81 (*smc-5(ok2421)*/mIn1[*dpy-10(e128)* mIs14] *II*; *unc-5(e791) IV*)

DLW131 (*smc-5(ok2421)*/mIn1[*dpy-10(e128)* mIs14] *II*; *lig-4(ok716) brc-1(xoe4) III*)

DLW134 (*smc-5(ok2421)*/mIn1[*dpy-10(e128)* mIs14] *II*; *polq-1(tm2572) brc-1(xoe4) III*)

DLW137 (*smc-5(ok2421)*/mIn1 [mIs14 *dpy-10(e128)*] *II*; *brc-1(xoe4) III*)

DLW157 (*brc-1(xoe4) III*; *unc-5(e791) IV*)

DLW175 (*smc-5(syb3065* [::AID*::3xFLAG]) II; *brc-1(xoe4) III*)

DLW182 (*smc-5(ok2421)*/mIn1[*dpy-10(e128)* mIs14] *II*; *GFP::brc-1 III*)

DLW202 (*smc-5(ok2421)*/mIn1 [*dpy-10(e128)* mIs14] *II*; *dpy-13(e184sd) unc-5(ox171::Mos1) IV*; KrIs14 [*phsp-16.48::MosTransposase*; *lin-15B?*; *punc-122::GFP*] *V*)

DLW203 (*brc-1(xoe4) III*; *dpy-13(e184sd) unc-5(ox171::Mos1) IV*; KrIs14 [*phsp-16.48::MosTransposase*; *lin-15B*; *punc-122::GFP*] *V*)

JEL515 (*GFP::brc-1 III*) JEL730 (*brc-1(xoe4) III*)

PHX3065 (*smc-5(syb3065* [::AID*::3xFLAG]*) II*)

YE57 (*smc-5(ok2421)*/mIn1 [mIs14 *dpy-10(e128)*] *II*)

Double and triple mutants which carried the *smc-5(ok2421)* and *brc-1(xoe4)* alleles incurred mutations within ∼6-10 generations of propagation, as indicated by progeny with movement defects, body morphology defects, or the presence of male offspring. To minimize the risk of *de novo* suppressor or enhancer mutations influencing the phenotypes we observed in these mutants, we froze stocks of these double and triple mutants at −80°C within 3 generations of the strains’ construction. All experiments using these strains were carried out on stocks which had been maintained for less than 1-2 months. If a strain began to segregate mutant phenotypes, a new isolate of the freshly generated strain was thawed from frozen stocks.

### CRISPR/Cas9 genome editing

CRISPR/Cas9 genome editing was performed by SUNY Biotech to generate the *smc-5(syb3065)* allele in which the endogenous sequence of *smc-5* is modified at its C terminus to code for both an AID* tag (peptide sequence PKDPAKPPAKAQVVGWPPVRSYRKNVMVSCKSSGGPEAAAFVK) and a 3xFLAG tag (peptide sequence DYKDHDGDYKDHDIDYKDDDDK). The coding sequence of *smc-5*, the AID* tag, and the 3xFLAG tag were respectively connected by flexible GAGS peptide linkers. The repair template for this insertion was synthesized as a single strand oligo and was injected with Cas9 enzyme and a single guide RNA targeting the 12^th^ exon of the *smc-5* locus. Successful integration was confirmed by PCR and Sanger sequencing. CRISPR edited strains were backcrossed three times to N2 before experiments were performed.

### *C. elegans* brood viability assays and Bayesian hierarchical modeling analysis

L4 stage hermaphrodite nematodes of each genotype to be scored were isolated 16-18hrs before irradiation was to be performed and were maintained at 15°C on NGM plates seeded with OP50. These worms were then exposed to 0, 2500, or 5000 Rads of ionizing radiation from a Cs^137^ source (University of Oregon). Following irradiation, n=5 hermaphrodites of each genotype and treatment combination were placed onto individual NGM plates seeded with OP50 and were maintained at 20°C. At 10hrs, 22hrs, and 46hrs post irradiation, the irradiated hermaphrodites were transferred to new NGM plates seeded with OP50. 58hrs after irradiation, the parent hermaphrodites were discarded. The proportion of F1 progeny which hatched, did not hatch (‘dead eggs’ indicating embryonic lethality), or were unfertilized on each plate was scored 36-48hrs after the removal of the parent hermaphrodite from a plate. The brood size of each hermaphrodite was calculated as (hatched progeny) + (dead eggs). Brood viability at each timepoint was calculated as (hatched progeny) / (brood size). Brood viability assays were performed in triplicate with the exception of *smc-5(syb3065)*, which was replicated twice with n=5 hermaphrodites scored for each radiation treatment in each replicate.

Brood viabilities of individual hermaphrodites for each given genotype and irradiation treatment were analyzed using RStan (Stan Development Team 2021). The brood viability data of individual hermaphrodites (h) for each genotype (g), timepoint scored (t), and irradiation treatment (i) was fit to a Beta-Binomial model:

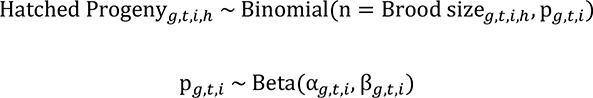

A metric (termed “gamma”) for the effect of ionizing radiation on the observed brood viability of each genotype was calculated in the Generated Quantities block during MCMC sampling from the posterior probability distribution of the parameter p, defined as:

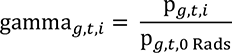

In addition to the model fit statistics output from Stan, model fit was assessed by posterior simulations. The expected brood viability for 1000 parent hermaphrodites from each genotype, timepoint, and irradiation treatment were simulated (Supplemental Figure 4B). For each simulated parent hermaphrodite, a brood size was sampled from the empirical data of the corresponding experimental group, values for α and β were sampled from the respective posterior probability distributions, and a value for p was simulated from a Beta distribution with shape parameters α and β. The number of hatching progeny were simulated ∼Binomial (brood size, p).

### Intersister/intrachromatid repair assay (ICR Assay)

ICR assays were performed as described in (Toraason *et al*. 2021a; b). Parent (P0) hermaphrodites for the ICR assay for each genotype were generated by crossing (see cross schemes detailed below).

### ICR assay cross schemes

1. Wild type (N2): P0 hermaphrodites were generated by crossing: (1) N2 males to DLW14 hermaphrodites to generate *unc-5(lib1)*/+ IV; KrIs14/+ V males; (2) F1 males to CB791 hermaphrodites to generate *unc-5(lib1)/unc-5(e791)* IV; KrIs14/+ V hermaphrodites.
2. *brc-1* mutant: P0 hermaphrodites were generated by crossing: (1) JEL730 males to DLW156 hermaphrodites to generate *brc-1(xoe4)* III; *unc-5(lib1)*/+ IV; KrIs14/+ V males; (2) F1 males to DLW157 hermaphrodites to generate *brc-1(xoe4)* III; *unc-5(lib1)*/*unc-5(e791)* IV; KrIs14/+ V hermaphrodites.
3. *smc-5* mutant: P0 hermaphrodites were generated by crossing: (1) YE57 males to DLW23 hermaphrodites to generate *smc-5(ok2421)*/mIn1 II; *unc-5(lib1)*/+ IV; KrIs14/+ V males; (2) F1 males to DLW81 hermaphrodites to generate *smc-5(ok2421)* II; *unc-5(lib1)*/*unc-5(e791)* IV; KrIs14/+ V hermaphrodites.

In brief, P0 hermaphrodites of the desired genotype were isolated 16-18hrs before heat shock and were maintained at 15°C. Heat shock was performed in an air incubator (refrigerated Peltier incubator, VWR Model VR16P) for one hour. The P0 worms were then allowed to recover at 20°C for nine hours. P0 hermaphrodites were placed onto individual NGM plates seeded with OP50 and maintained at 20°C. 22hrs, 34hrs, and 46hrs after heat shock, the P0 worms were transferred to new NGM plates seeded with OP50. 58hrs after heat shock, P0 hermaphrodites were removed from their NGM plates and discarded. Plates with P0 hermaphrodites were maintained at 20°C, while plates with F1 progeny were placed at 15°C.

F1 progeny were scored for GFP fluorescence ∼54-70hrs after the P0 hermaphrodite was removed. ∼18hrs before scoring, plates with F1 progeny were placed at 25°C to enhance GFP expression. Fluorescent phenotype scoring was performed on a Axio Zoom v16 fluorescence stereoscope (Zeiss). F1 progeny which expressed recombinant fluorescence phenotypes were isolated and lysed for sequencing (see Sequencing and analysis of ICR assay conversion tracts). Nonrecombinant progeny were discarded. If all progeny on a plate were in larval developmental stages at the time of scoring, then the number of dead eggs and unfertilized oocytes were additionally recorded.

ICR assays in *brc-1(xoe4)* and *smc-5(ok2421)* mutants were replicated 4 times and the broods of at least 20 parent hermaphrodites scored in each replicate. The ICR assay in a wild type genetic background was performed once and combined with previous data (Toraason *et al*. 2021a).

We observed increased GFP+ progeny in the ICR assays we performed in both *brc-1* and *smc-5* mutant backgrounds as compared to wild type (Supplemental Figure 1). This result was unexpected, as the ICR assay is performed in parent hermaphrodites which are heterozygous for the ICR assay construct and an allele of *unc-5* which does not carry any GFP homology. Thus, the homolog is not a viable repair template to restore GFP fluorescence and we would expect that DSB repair should be ultimately directed towards intersister/intrachromatid repair templates regardless of the genetic background. This increased proportion of GFP+ progeny in *brc-1* and *smc-5* mutants may indicate altered bias for the upstream intersister/intrachromatid nonallelic GFP repair template as compared to the allelic repair template. Allelic recombination in the ICR assay reincorporates the Mos1 transposon into the final repair product and therefore does not yield a detectable event, so a reduced propensity for this template engagement would increase the number of GFP+ recombination events we identify. The tandem GFP sequences of the ICR assay contain polymorphisms (Toraason *et al*. 2021a), and the presence of nucleotide polymorphisms between damaged DNA sequences and recombination repair templates is known to reduce the likelihood of recombination between loci (Chen and Jinks-Robertson 1999; Hum and Jinks-Robertson 2019). It is therefore possible that BRC-1 and SMC-5/6 play some role either in the detection of polymorphisms during the strand invasion step of recombination or in facilitating the rejection of heteroduplex recombination intermediates. Previous work has shown that BRC-1 restricts heterologous recombination (León-Ortiz *et al*. 2018), consistent with a role for BRC-1 in rejecting repair templates with sequence divergence.

Alternately, the elevated rate of GFP+ progeny we observed may be the product of increased Mos1 mobilization in the germlines of *brc-1* and *smc-5* mutants. We propose that this is a less likely explanation for the rates of GFP+ progeny in the *brc-1* and *smc-5* ICR assays, as the frequencies of non-Unc progeny were not broadly elevated in the *brc-1* and *smc-5* interhomolog assays, which assess Mos1 excision at the same locus as the ICR assay using an identical Mos1 transposase transgene construct (Supplemental Figure 2). As we cannot specifically delineate the underlying mechanisms which increase the rates of GFP+ progeny in *brc-1* and *smc-5* mutants, the frequency of ICR assay recombinants in this study should not necessarily be extrapolated to represent an absolute increase in rates of intersister/intrachromatid recombination more broadly in these contexts.

### Sequencing and analysis of ICR assay conversion tracts

Recombinant ICR assay progeny were placed in 10μL of 1x Worm Lysis Buffer for lysis (50mM KCl, 100mM TricHCl pH 8.2, 2.5mM MgCl_2_, 0.45% IGEPAL, 0.45% Tween20, 0.3μg/μL proteinase K in ddH_2_O) and were iteratively frozen and thawed three times in a dry ice and 95% EtOH bath and a 65°C water bath. Samples were then incubated at 60°C for one hour and 95°C for 15 minutes to inactive the proteinase K. Final lysates were diluted with 10μL ddH2O.

Conversion tracts were PCR amplified using OneTaq 2x Master Mix (New England Biolabs). Noncrossover recombination products were amplified using forward primer DLO822 (5’-ATTTTAACCCTTCGGGGTACG-3’) and reverse primer DLO823 (5’-TCCATGCCATGTGTTAATCCCA-3’). Crossover recombination products were amplified using forward primer DLO824 (5’-AGATCCATCTAGAAATGCCGGT-3’) and reverse primer DLO546 (5’-AGTTGGTAATGGTAGCGACC-3’). PCR products were run on an Agarose gel and desired bands were isolated by gel extraction (QIAquick Gel Extraction Kit, New England Biolabs) and were eluted in ddH_2_O. Amplicons were submitted for Sanger sequencing (Sequetech) with three primers.

Noncrossovers were sequenced using DLO822, DLO823, and DLO1077 (5’-CACGGAACAGGTAGGTTTTCCA-3’) and crossovers were sequenced using DLO824, DLO546, and DLO1077.

Sanger sequencing chromatograms were analyzed using Benchling alignment software (Benchling) to determine converted polymorphisms. Heteroduplex DNA signals were identified by two prominent peaks in the chromatogram at the site of a known polymorphism. Putative heteroduplexed samples were PCR amplified and submitted for sequencing a second time for confirmation as described above.

Samples which produced PCR products of the expected size but did not yield interpretable sequencing were subsequently analyzed using TOPO cloned amplicons. ICR assay locus amplicons were PCR amplified as described above but were immediately cloned into pCR2.1 vector using the Original TOPO-TA^TM^ Cloning Kit^TM^ (Invitrogen) following kit instructions. Putative successful amplicon clones were identified by PCR amplification using 2xOneTaq Master Mix (New England Biolabs) with primers DLO883 (5’-CAGGAAACAGCTATGACCATG-3’) and DLO884 (5’-TGTTAAAACGACGGCCAGGT-3’). Plasmids containing amplicon inserts were isolated from 2mL LB+Amp cultures using the GENEJET Miniprep kit (Fischer Scientific) and were submitted for Sanger sequencing (Sequetech) using primers DLO883 and DLO884.

To acquire additional wild type ICR assay crossover tracts for our analyses, three “bulk” replicates of the wild type ICR assay were performed following the protocol described in the ‘Intersister/intrachromatid repair assay’ with the following exceptions: 1) n=3 hermaphrodites were passaged together on individual plates during the experiment; 2) transfers were only performed at 10hr, 22hr, and 46 hr following heat shock; and, 3) plates were screened for body wall GFP+ crossover recombinants but the frequency of pharynx GFP+ and GFP-nonrecombinant progeny were not scored. Body wall GFP+ crossover progeny were lysed and following the preceeding protocol.

Not all lysed recombinant yielded successful PCR products or sequences. Of the additional wild type ICR assay recombinants sequenced for this manuscript, 11 of 11 noncrossover and 52 of 52 crossover lysates were successfully sequenced. Among lysates from *brc-1* mutant ICR assays, 37 of 37 noncrossover and 70 of 73 crossover lysates were successfully sequenced. Among lysates from *smc-5* mutant ICR assays, 56 of 56 noncrossover and 27 of 28 crossover lysates were successfully sequenced.

### Interhomolog assay (IH assay)

IH assays were performed as described in (Rosu *et al*. 2011). In brief, P0 hermaphrodites were generated by crossing (see cross schemes detailed below).

IH assay cross schemes:

1. Wild type (N2): P0 hermaphrodites were generated by crossing: (1) N2 males to AV554 hermaphrodites to generate *dpy-13(e184sd) unc-5(ox171::Mos1)*/+ IV; KrIs14/+ V males; (2) F1 males to CB791 hermaphrodites to generate *dpy-13(e184sd) unc-5(ox171::Mos1)*/*unc-5(e791)* IV; KrIs14/+ V hermaphrodites.
2. *brc-1* mutant: P0 hermaphrodites were generated by crossing: (1) JEL730 males to DLW203 hermaphrodites to generate *brc-1(xoe4)* III; *dpy-13(e184sd) unc-5(ox171::Mos1)*/+ IV; KrIs14/+ V males; (2) F1 males to DLW157 hermaphrodites to generate *brc-1(xoe4)* III; *dpy-13(e184sd) unc-5(ox171::Mos1)*/*unc-5(e791)* IV; KrIs14/+ V hermaphrodites.
3. *smc-5* mutant: P0 hermaphrodites were generated by crossing: (1) YE57 males to DLW23 hermaphrodites to generate *smc-5(ok2421)*/mIn1 II; *dpy-13(e184sd) unc-5(ox171::Mos1)*/+ IV; KrIs14/+ V males; (2) F1 males to DLW81 hermaphrodites to generate *smc-5(ok2421)* II; *dpy-13(e184sd) unc-5(ox171::Mos1)*/*unc-5(e791)* IV; KrIs14/+ V hermaphrodites.

The heat shock and timing at which parent hermaphrodites were transferred to new NGM plates was performed identically to the ICR assay (see ‘Intersister/Intrachromatid repair assay (ICR assay)’ above). However, the number of eggs and unfertilized oocytes laid by each hermaphrodite was recorded immediately following the removal of the parent hermaphrodite at each timepoint and plates carrying F1 progeny were maintained at 20°C. Plates were scored for F1 wild type moving (non-Unc) progeny ∼84-96hrs after parent hermaphrodites were removed. F1 Unc progeny were discarded.

F1 non-Unc progeny were placed on single NGM plates seeded with OP50 bacteria. Dpy non-Unc progeny (putative noncrossover recombinants) were lysed following the protocol described in ‘Sequencing and analysis of SCR assay conversion tracts’. If Dpy non-Unc progeny died before the time of lysis and had laid F2 progeny, non-Unc segregant F2s were lysed instead. Non-Dpy non-Unc progeny (putative crossover recombinants) were allowed to lay F2 progeny. If progeny were laid and Dpy non-Unc F2 segregants were identified, these Dpy non-Unc F2s were lysed and the F1 was inferred not to be a crossover recombinant. If >50 F2 progeny were on the plate and no Dpy non-Unc segregants were identified, the F1 was assumed to be a crossover recombinant and no worms were lysed. If very few progeny were laid and no Dpy non-Unc segregants were identified, the F1 non-Unc or its non-Unc F2 offspring were lysed and subsequently subjected to PCR genotyping analysis using OneTaq 2x Master Mix (New England Biolabs) to determine the genotype of *unc-5* and *dpy-13*. The presence of Mos1 in the *unc-5* locus was assessed using primers DLO987 (5’-TCTTCTTGCCAAAGCGATTC-3’) and DLO1082 (5’-TTCTCTCCGAGCAATGTTCC-3’). The *dpy-13* locus was assessed using primers DLO151 (5’-ATTCCGGATGCGAGGGAT-3’) and DLO152 (5’-TCTCCTCGCAAGGCTTCTGT-3’). Lysed F1 nUnc nDpy progeny were inferred to be crossover recombinants if the worms 1) carried the Mos1 transposon at the *unc-5* locus and were heterozygous for the *dpy-13(e184)* allele, or; 2) did not carry the Mos1 transposon at the *unc-5* locus and were homozygous wild type for *dpy-13*.

The *unc-5* locus was amplified for sequencing by PCR using OneTaq 2x Master Mix (NEB) with primers DLO1081 (5’-TCTTTTCAGGCTTTGGCACTG-3’) and DLO1082. PCR products were run on an agarose gel and desired bands were isolated by gel extraction (QIAquick Gel Extraction Kit, New England Biolabs) and were eluted in ddH_2_O. These amplicons were submitted for Sanger sequencing (Sequetech) with primer DLO1082 or DLO150 (5’-GTTCCATGTTTGATGCTCCAAAAG-3’). Sanger sequencing chromatograms were compared to the wild type *unc-5* sequence using Benchling alignment software. Samples which showed a reversion to wild-type *unc-5* sequence at the site of Mos1 excision were inferred to be noncrossover recombinants. Samples which showed mutations that preserved the reading frame of the *unc-5* locus were considered ‘mutant non-Unc’. One of the five *brc-1* IH assay mutant non-Uncs we sequenced carried two distinguishable mutagenic repair products. These two mutations likely represent the outcomes of both a meiotic DSB repair event and an additional somatic repair event in the progeny. We have previously observed analogous somatic Mos1 excision events in F1 progeny in the ICR assay (Toraason *et al*. 2021a; b). As we cannot distinguish the source of the respective repair events, this mutant was excluded from subsequent sequence analysis (Supplemental Figure 3).

Samples which showed mixed sequences despite a clear amplicon being generated in the PCR were subsequently TOPO cloned, as described in ‘Sequencing and analysis of ICR assay conversion tracts’, except that the amplicon used in the reaction was generated using primers DLO1081 and DLO1082.

Not all interhomolog assay non-Unc progeny were able to be confirmed as recombinants by sequencing. Of the wild type IH assay non-Unc progeny identified, 176 of 178 putative noncrossovers were successfully sequenced. Among lysates from *brc-1* mutant IH assays, 72 of 76 putative noncrossovers were successfully sequenced. Among lysates from *smc-5* mutant IH assays, 213 of 229 putative noncrossovers were successfully sequenced. Non-Unc progeny whose *unc-5* DNA repair events could not be identified by sequencing were considered ‘undetermined non-Unc’ in subsequent analyses of this data.

### EdU Sister Chromatid Exchange Assay

EdU Sister Chromatid Exchange assays were performed as described in (Almanzar *et al*. 2021, 2022).

### Immunofluorescence localization of SMC-5/6 and BRC-1

Immunofluorescence was performed as in (Libuda *et al*. 2013) or (Howe *et al*. 2001). For both protocols, L4 staged hermaphrodites were isolated 18-22hrs before dissection and maintained at 20°C on NGM plates seeded with OP50. Nematodes which were irradiated preceding an immunofluorescence experiment were exposed to a Cs^137^ source (University of Oregon) were dissected less than an hour after following irradiation. Samples prepared for GFP::BRC-1 visualization were dissected in 1x Egg Buffer (118 mM NaCl, 48 mM KCl_2_, 2 mM CaCl_2_, 2 mM MgCl_2_, 25 mM HEPES pH7.4, 0.1% Tween20) and were fixed in 1x Egg Buffer with 1% paraformaldehyde for 5 min on a Superfrost Plus slide (VWR). Samples prepared for SMC-5::AID*::3xFLAG visualization were dissected in 1x Sperm Salts (50mM PIPES pH7, 25mM KCl, 1mM MgSO_4_, 45mM NaCl, 2mM CaCl_2_) and an equal volume of 1x Sperm Salts with 3% paraformaldehyde was applied for 5 min before samples were affixed to a Superfrost Plus slide (VWR). For both protocols, gonads were then flash frozen in liquid nitrogen and the cover slip was removed. Germlines stained for GFP::BRC-1 were then fixed for 1 min in ice cold MeOH and then were washed in 1x PBST (1x PBS, 0.1% Tween20), while germlines stained for SMC-5::AID*::3xFLAG were fixed for 1 min in ice cold 95% EtOH and then were washed in 1xPBST* (1x PBS, 0.5% Triton-X100, 1mM EDTA pH8). Slides were then washed 3x in PBST or PBST* respectively before being placed in Block (1xPBST or 1xPBST* with 0.7% bovine serum albumin) for at least 1 hour.

Primary antibody staining was performed by placing 50μL of antibody diluted in PBST for samples in which GFP::BRC-1 or RAD-51 were to be visualized or PBST* if the sample was to be stained for SMC-5::AID*::3xFLAG (see below for specific dilutions of primary antibodies). A parafilm coverslip was placed on each sample and the slides were incubated for 16-18hrs in a dark humidifying chamber. Slides were then washed 3x in PBST or PBST* for 10 min. 50μL of secondary antibody diluted in PBST for samples in which GFP::BRC-1 or RAD-51 were to be visualized or PBST* if the sample was to be stained for SMC-5::AID*::3xFLAG (see below for specific dilutions of primary antibodies) was then placed on each slide. Slides were incubated for 2hrs in a dark humidifying chamber with a parafilm coverslip. Slides were then washed 3x in PBST or PBST* for 10 min in a dark chamber. 50μL of 2μg/mL DAPI was then applied to each slide. Slides were incubated in a dark humidifying chamber with parafilm coverslips for 5 min.

Slides were then washed 1x in PBST or PBST* for 5 min in a dark chamber before being mounted in VectaShield with a No. 1.5 coverslip (VWR) and sealed with nail polish. Slides were maintained at 4°C until imaging. All slides stained for SMC-5::AID*::3xFLAG were imaged within 48 hours of mounting. Immunofluorescence images were acquired at 512×512 pixel dimensions on an Applied Precision DeltaVision microscope. All images were acquired in 3D using Z-stacks at 0.2μm intervals and were deconvolved with Applied Precision softWoRx deconvolution software. Individual images of whole germlines were stitched as 3D Z-stacks in FIJI using the Grid/Collection Stitching plugin (Preibisch *et al*. 2009) or as maximum intensity projections using Photoshop (Adobe). The intensity levels of images displayed in this manuscript were adjusted in Photoshop for clarity.

The following primary antibodies were used In this study at the given dilutions: Chicken αRAD-51 (1:1000; (Kurhanewicz *et al*. 2020)), Mouse αmini-AID M214-3 (1:500, MBL International), Rat αHTP-3 (1:1000, this study), Rabbit αGFP (1:500 (Yokoo *et al*. 2012)).

Immunofluorescence staining of SMC-5/6 was further attempted using previously published antibodies (Bickel *et al*. 2010). We were unable to generate samples with specific staining using these antibodies, potentially due to their age. We additionally attempted to raise new antibodies using the previously published epitopes (Bickel *et al*. 2010) (synthesized by GenScript) in chickens (SMC-5) or rabbits (SMC-6). Neither of these antibodies exhibited specific staining.

### Antibody Generation

The HTP-3 antibody used in this study was generated from an identical C-terminal segment of the HTP-3 protein (synthesized by GenScript) as was used by (MacQueen *et al*. 2005). Antibodies were produced in rats and affinity purified by Pocono Rabbit Farms.

### Statistics

All statistics were calculated in R (v4.0.3). Specific tests utilized are described in text or in the figure legends. Data wrangling was performed using the Tidyverse (v1.3.0). Bayesian hierarchical models were fit using Rstan (v2.21.2). Binomial Confidence Intervals were calculated using the DescTools package (v 0.99.38).

### Data Availability

All data generated or analyzed in this study are included in the manuscript and supporting files. Source data files have been provided for Figures 1A, 1C, 1D, 1E, 2A, 2B, 2D, 2E, 3 (all panels), and 4A. Source data files have also been provided for Supplemental Figures 1 (all panels), 2 (all panels), 4A, and 6A. Source code files have been provided for Figure 4A.

---

**Supplemental Figure 1.**
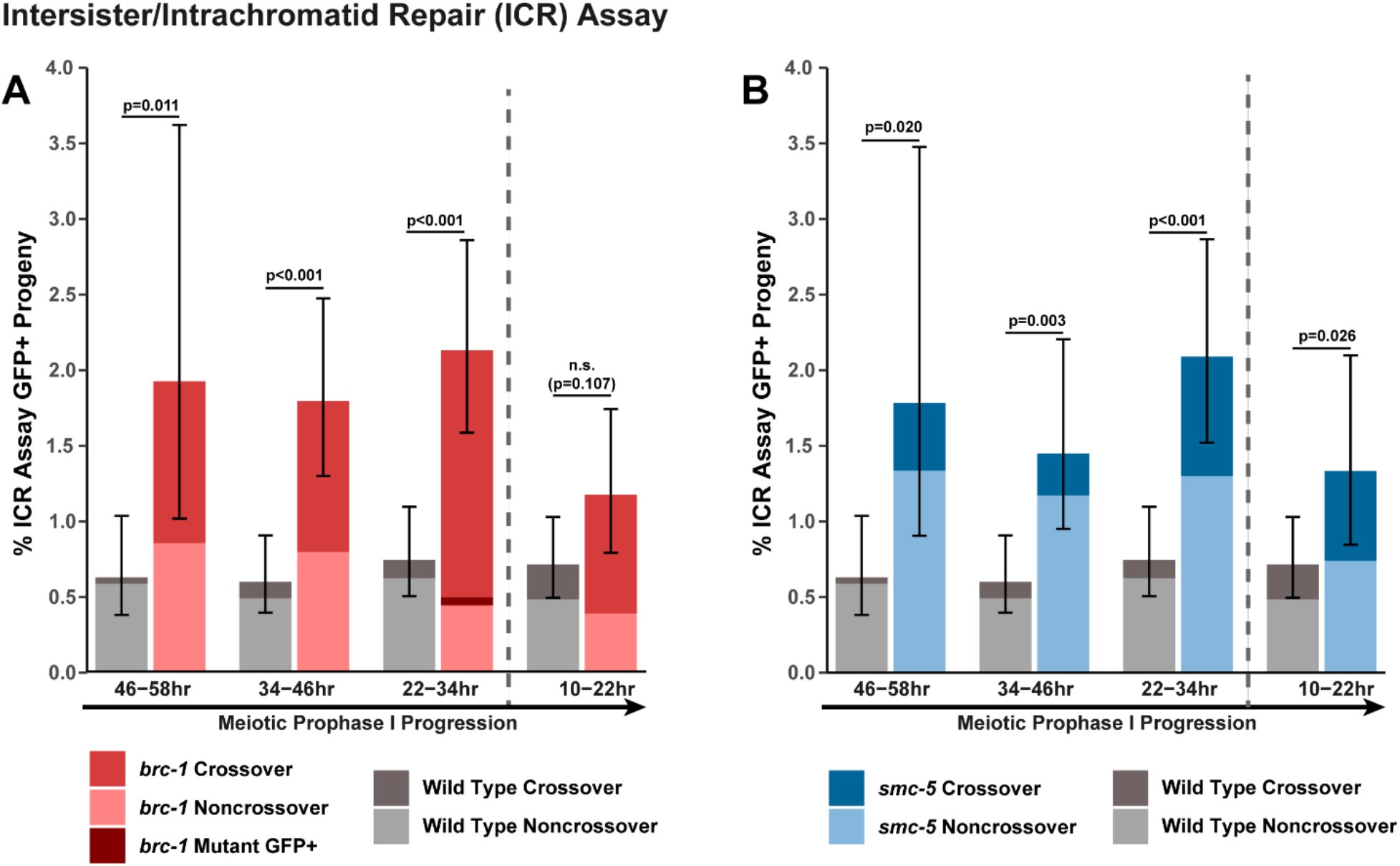
Intersister/intrachromatid repair (ICR) assay GFP+ progeny are elevated in *brc-1* and *smc-5* mutants. Stacked bar plots displaying the percent of all progeny scored in wild type and *brc-1* (A) or *smc-5* (B) ICR assays which were determined to be GFP+ noncrossover recombinants, crossover recombinants, or mutants. Error bars represent the 95% Binomial confidence intervals for the frequencies of GFP+ progeny. P values were calculated by Fisher’s Exact test. Vertical dashed lines demarcate the interhomolog window (22-58hr post heat shock) and non-interhomolog window (10-22hr post heat shock) timepoints. **Supplemental Figure 1 – source data 1. The source data for** Supplemental Figure 1 **is provided.** [Supplemental Figure 1 source data.xlsx]. The total number of ICR assay progeny with GFP+ or non-GFP+ phenotypes are listed. Wild type data is shared with Figure 1 and Figure 2.

**Supplemental Figure 2.**
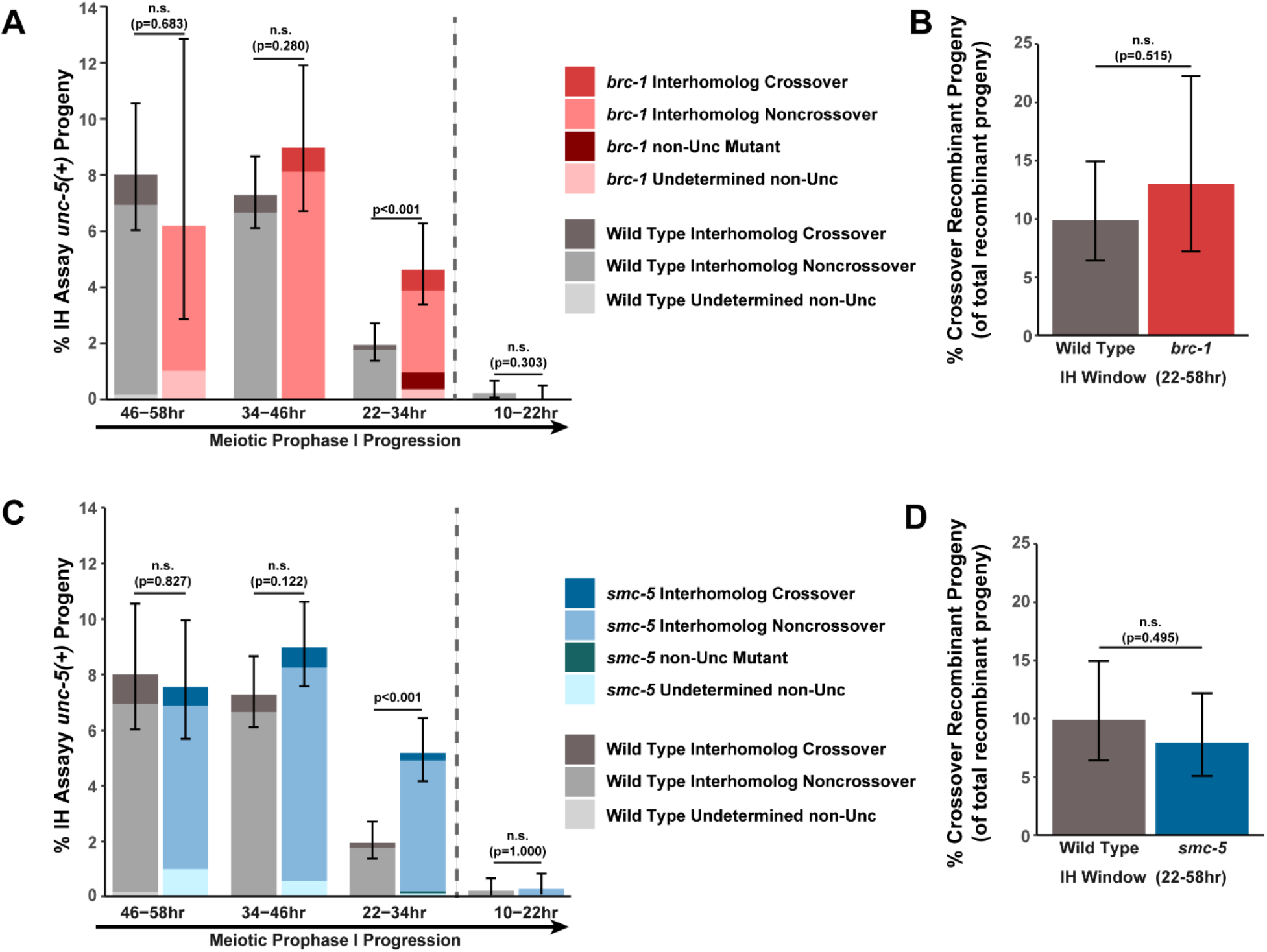
Interhomolog repair is largely unperturbed in *brc-1* and *smc-5* mutants. A) Stacked bar plots displaying the percent of all progeny scored in wild type and *brc-1* IH assays which were determined to be noncrossover recombinants, crossover recombinants, non-Unc mutants, or undetermined non-Unc. B) Percent of all recombinant progeny identified within the interhomolog window of wild type and *brc-1* IH assays which were crossover recombinants. C) Stacked bar plots displaying the percent of all progeny scored in wild type and *smc-5* IH assays which were determined to be noncrossover recombinants, crossover recombinants, non-Unc mutants, or undetermined non-Unc. D) Percent of all recombinant progeny identified within the interhomolog window of wild type and *smc-5* IH assays which were crossover recombinants. Error bars represent the 95% Binomial confidence intervals for the frequencies of non-Unc progeny. P values were calculated by Fisher’s Exact test. Vertical dashed lines demarcate the interhomolog window (22-58hr post heat shock) and non-interhomolog window (10-22hr post heat shock) timepoints. **Supplemental Figure 2 – source data 1. The source data for** Supplemental Figure 2 **is provided.** [Supplemental Figure 2 source data 1.xlsx]. The total number of IH assay progeny with recombinant or mutant nonUnc phenotypes or Unc nonrecombinant phenotypes are listed. Wild type data is shared with Figure 1 and Figure 2.

**Supplemental Figure 3.**
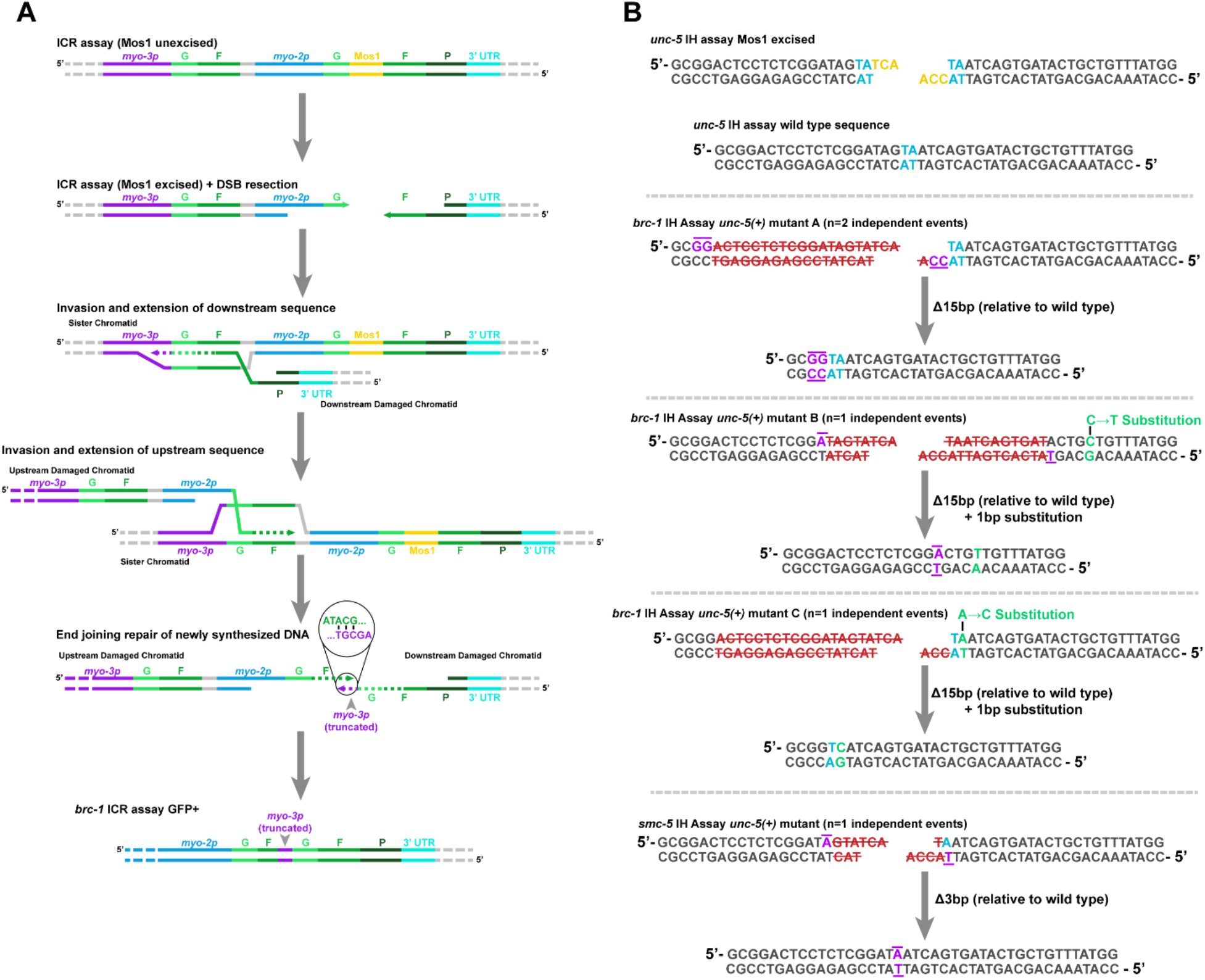
Illustrations of mutants identified in ICR and IH assays. A) Illustrated depiction of ICR assay GFP+ mutant identified in a *brc-1* mutant background (Figure 1D, Supplemental Figure 1A). The partial tandem duplication produced (bottom) can best be parsimoniously explained by two independent strand invasion and extension events on either end of the DSB. For simplicity, intersister recombination is depicted in this diagram. However, intrachromatid templates could also have been engaged to produce the final product. B) Illustrations of *unc-5* lesions identified in IH assay non-Unc progeny in *brc-1* or *smc-5* mutants. Specific mutation signatures are separated by horizontal dashed grey lines. The wild type *unc-5* locus sequence at the site of Mos1 excision and the DSB product generated by Mos1 excision are displayed on the top of panel B. Blue letters indicate a duplicated TA at the site of Mos1 insertion in the *unc-5(ox171)* locus, while yellow letters indicate the 3nt 3’ overhangs left following Mos1 excision (Robert *et al*. 2008). In the panels displaying mutations identified, purple letters with bars indicate complementary bases flanking the deletion site. Red letters struck through with red lines indicate bases in the damaged locus which were deleted to produce the final product. Green letters indicate sites of nucleotide substitution mutations.

**Supplemental Figure 4.**
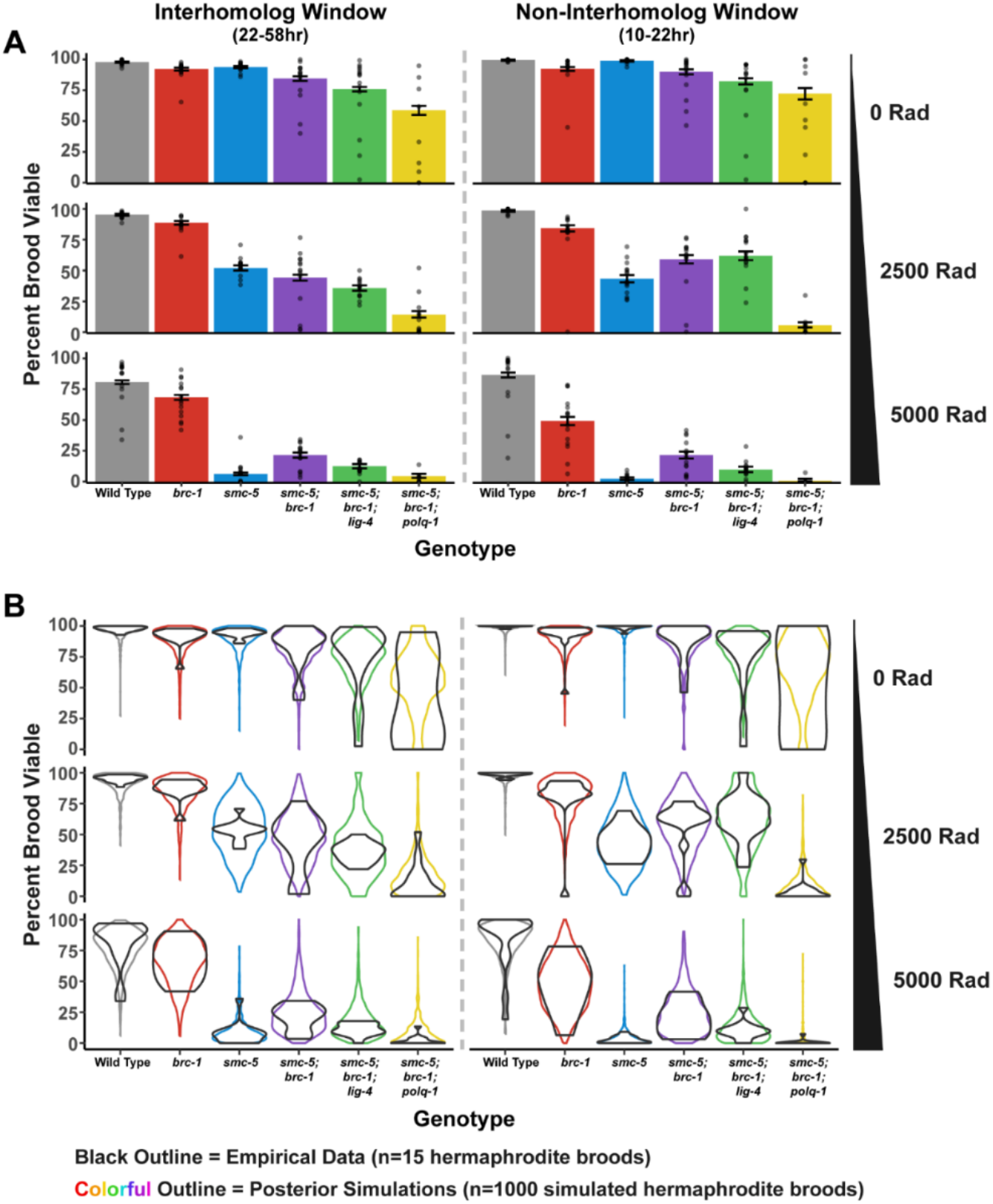
Brood viability results following irradiation. A) Brood viability results following irradiation at doses of 0, 2500, or 5000 Rads. Bars represent the population brood viability, while points represent the brood viabilities of individual hermaphrodites scored. Error bars indicate 95% Binomial confidence intervals of the population brood viability. B) Violin plots of empirical brood viabilities from individual hermaphrodites scored (displayed as points in A) and posterior simulations from the Beta-Binomial model fit to the data (Figure 4A, see Methods). In all panels, vertical dashed grey lines separate interhomolog (22-58hr post heat shock) and non-interhomolog window (10-22hr post heat shock) timepoints. **Supplemental Figure 4 source data 1. The source data for** Supplemental Figure 4 **is provided.** [Supplemental Figure 4 source data 1.xlsx]. The number of hatched (Live), unhatched (Dead), or unfertilized (Unf) F1 progeny scored in the brood viability experiment data used to generate Figure 1. The number of progeny scored are separated by individual timepoints (Timept) for each parent scored (Plate_ID). Experimental replicates are delinated by the date of irradiation treatment (IR_date). Wild type and *smc-5(ok2421)* data is shared with Supplemental Figure 6.

**Supplemental Figure 5.**
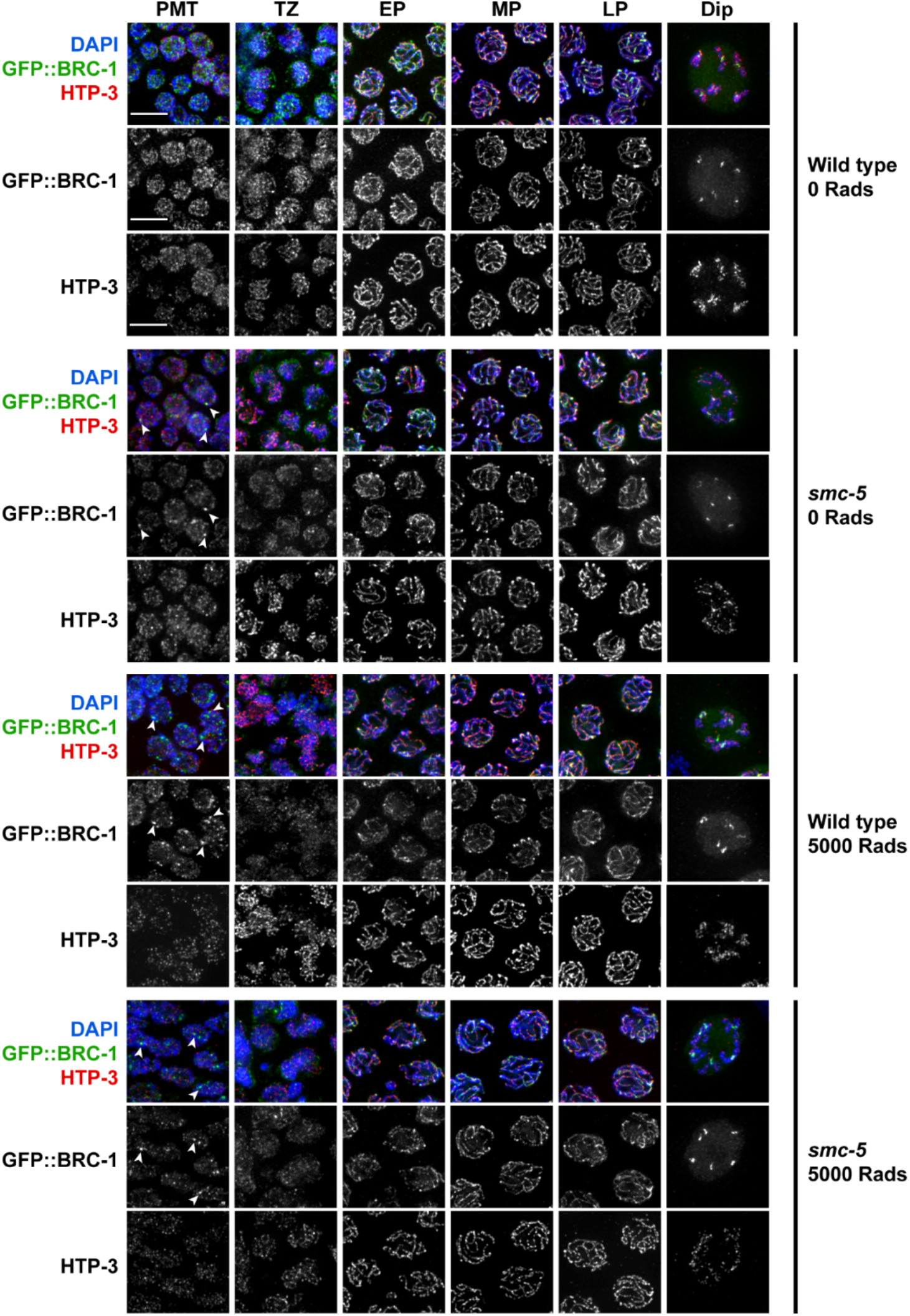
SMC-5/6 is not required for GFP::BRC-1 localization. Deconvolved widefield images of germline nuclei stained for GFP (GFP::BRC-1), chromosome axis protein HTP-3, and DAPI (DNA) in a wild type or *smc-5(ok2421)* mutant background and treated with 0 or 5000 Rads of ionizing radiation. Scale bars represent 5μm. Stages of meiotic nuclei were determined based on DAPI morphology and are listed on the top of the figure (PMT = premeiotic tip, TZ = transition zone, EP = early pachytene, MP = mid pachytene, LP = late pachytene, Dip = Diplotene). Arrowheads indicate GFP::BRC-1 foci.

**Supplemental Figure 6.**
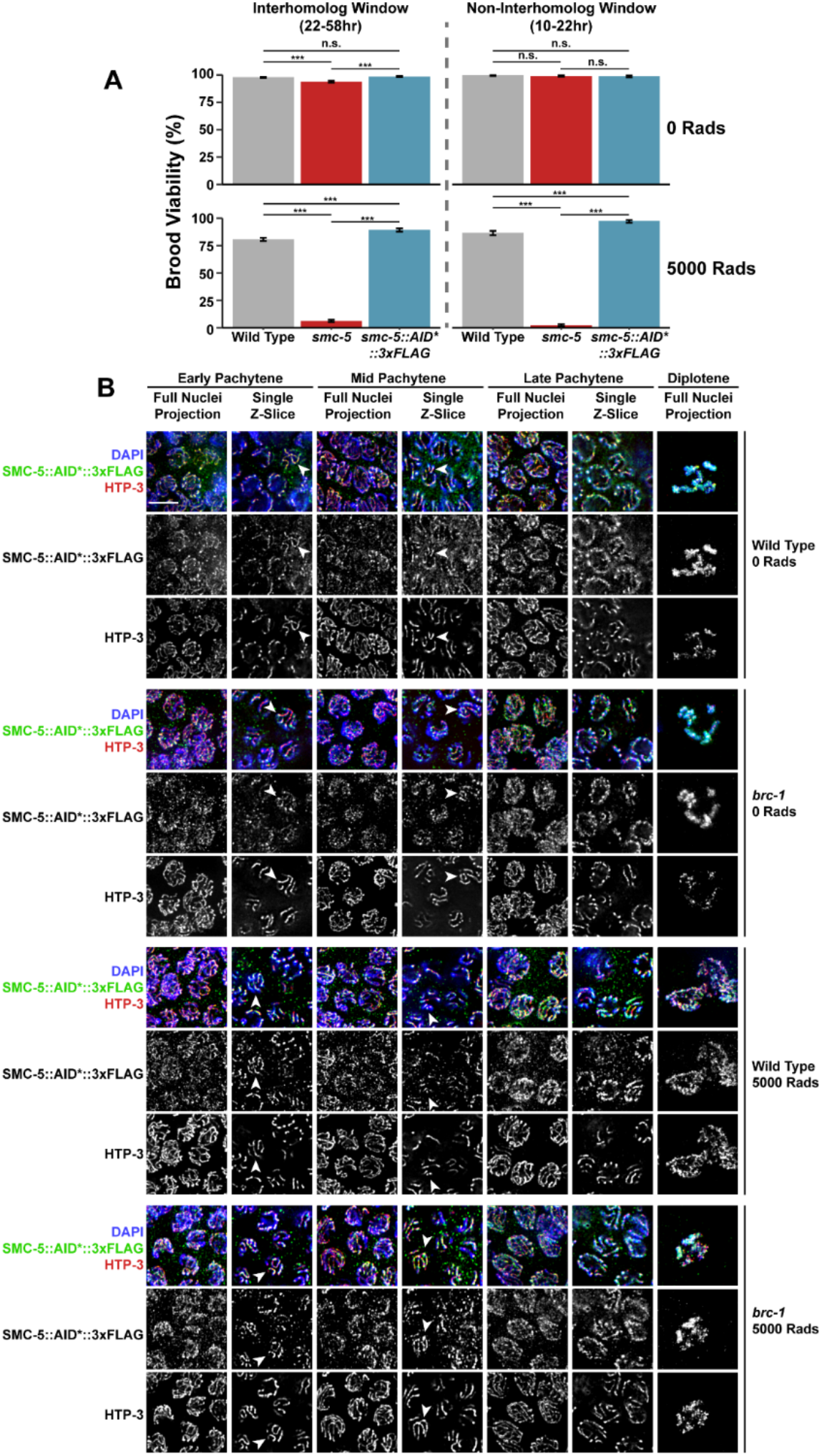
BRC-1 is not required for SMC-5::AID*::3xFLAG localization. A) Brood viability of wild type, *smc-5(ok2421)*, and *smc-5(syb3065)* hermaphrodites exposed to 0 or 5000 Rads of ionizing radiation. Bars represent the population brood viability of each strain. P values were calculated by Fisher’s Exact Test (n.s. = not significant p>0.05, *** p<0.001). Error bars represent the 95% Binomial confidence interval of the brood viability estimate. B) Deconvolved images of germline nuclei stained for AID* (SMC-5::AID*::3xFLAG), chromosome axis protein HTP-3, or DAPI (DNA) in a wild type or *brc-1(xoe4)* mutant background and treated with 0 or 5000 Rads of ionizing radiation. Scale bars represent 5μm. Stages of meiotic nuclei are determined based on DAPI morphology and are listed at the top of the figure. For each image, a max intensity projection of whole nuclei and single z-slices are displayed to demonstrate the relative localization of SMC-5 and HTP-3. Arrowheads indicate examples of colocalization between HTP-3 and SMC-5::AID*3xFLAG. **Supplemental Figure 6 source data 1. The source data for** Supplemental Figure 6A **is provided.** [Supplemental Figure 6 source data 1.xlsx]. The number of hatched (Live), unhatched (Dead), or unfertilized (Unf) F1 progeny scored in the brood viability experiment data used to generate Figure 1. The number of progeny scored are separated by individual timepoints (Timept) for each parent scored (Plate_ID). Experimental replicates are delineated by the date of irradiation treatment (IR_date). Wild type and *smc-5(ok2421)* data is shared with Supplemental Figure 4.

**Supplemental Figure 7.**
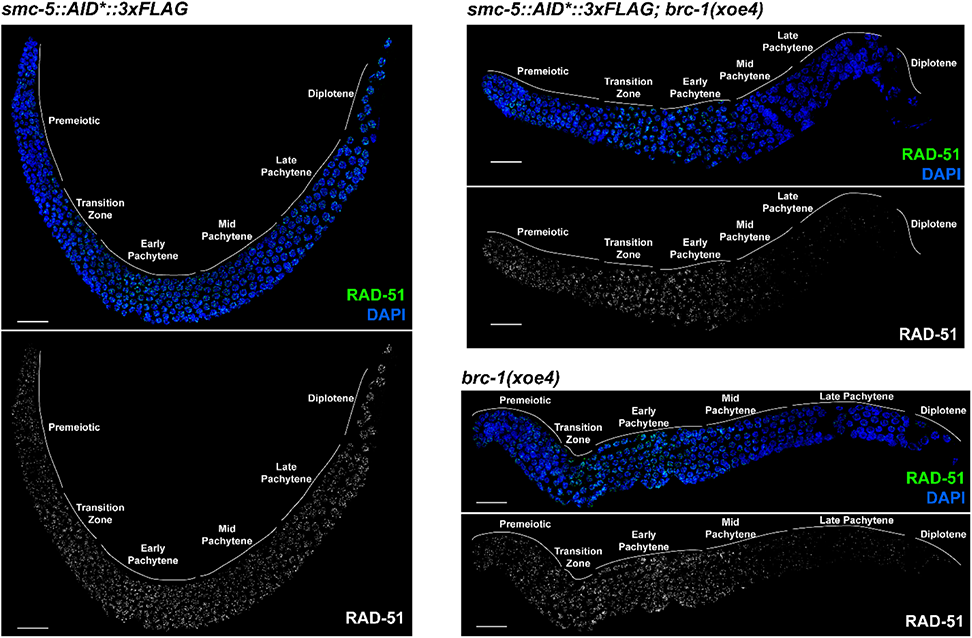
SMC-5::AID*::3xFLAG does not inhibit RAD-51 localization to irradiation-induced DSBs. Deconvolved images of whole extruded germlines stained for RAD-51 and DAPI. All germlines were exposed to 5000 Rads of ionizing radiation and were dissected within 1 hour of the radiation treatment. Loss of *brc-1* impedes RAD-51 localization in mid/late pachytene (Janisiw *et al*. 2018; Li *et al*. 2018), and this phenotype is not recapitulated nor enhanced by the *smc-5(syb3065)* allele. Grey lines and labels demarcate the mitotic and meiotic stages of the germline. Scale bars represent 20μm.

